# Geographic Variation and Host Genetics Shape the Human Skin Microbiome

**DOI:** 10.1101/2025.05.01.651599

**Authors:** Hoon Je Seong, Christopher Quince

## Abstract

The human skin microbiome is shaped by a complex interplay of host physiology, environmental exposure, and microbial interactions across domains of life. However, the relative contributions of host genetics and geography remain unresolved. Here, we present the first application of ultra-low coverage human genome imputation from skin metagenomic data, analysing 1,756 samples from multiple skin types and timepoints, with matched genotypes from 327 individuals across five countries. We also generate expanded Skin Microbial Genome Collection (eSMGC), comprising 675 prokaryotic, 12 fungal, 2,344 viral, and 4,935 plasmid genomes, correcting extensive false positives in existing references. Intercountry comparisons reveal that geography explains more microbiome variation than skin type, and that host genetics contributes previously uncharacterized structure—exemplified by distinct profiles in Chinese individuals. Genome-wide association analysis identifies 107 SNPs linked to 22 microbial taxa, including phages and plasmids, implicating host genes in skin structure, immunity, and lipid metabolism. *Cutibacterium acnes* and its phages exhibit geographic divergence and phage–host co-adaptation. Finally, host-infecting viruses, particularly papillomaviruses, are associated with elevated microbial diversity and immune-modulatory functions. These findings establish host genetics as a determinant of skin microbiome ecology and highlight the value of multi-domain, geographically diverse analyses.

## Introduction

The human skin serves as the outermost physical barrier of the body and hosts a wide variety of microorganisms, encompassing all domains of life^1^. These commensal microorganisms are intimately linked to the host’s skin condition, with the microenvironment^2^ and immune system^3^ combining to shape the structure of the skin microbiome. Previous studies have shown that the skin microbiome varies between skin types, e.g. sebaceous or dry, within an individual but also varies stably between individuals, even at the level of particular strains of *Cutibacterium acnes*^4,5^. The skin microbiome is also influenced by exogenous environmental factors, such as the degree of urbanization of the host environment^6^ and air pollution^7^. Interestingly, two distinct community clusters or cutotypes have been identified^8^ but only in Chinese individuals. This implies country of origin impacts the skin microbiome. A conclusion supported by further studies, but the relative importance of geography or host genetics in driving this difference has yet to be resolved^8,9^.

There have been very few studies on the association between human genetics and the skin microbiome and these have all used 16S rRNA amplicon sequencing, which can only provide prokaryotic community structure at a relatively low level of resolution^10,11^. Shotgun metagenomics, where all the DNA in a sample is sequenced^12^, captures all domains of life, prokaryotes, eukaryotes and phages to strain resolution but it is relatively challenging to apply to the skin because host DNA is often a large fraction of the resulting sample. Innovations in modern genomics have introduced ultra-low coverage imputation methods, enabling the accurate study of human genetics—even from ancient samples with minimal DNA^13^—with greater accuracy than traditional single nucleotide polymorphism (SNP) arrays^13^. This provides a unique opportunity, which we seize here, to utilize existing legacy skin metagenome samples, which span multiple countries, to disentangle the importance of geography and host genetics, across all domains of life at high resolution.

To characterize the skin microbiota we will use genome binning techniques that allow the multi-domain recovery of metagenome-assembled genomes (MAGs); these have filled gaps in our understanding of host^14^ and environmental microbiomes^15^, especially in ecosystems where microbial reference genomes are incomplete. They have also revealed the critical roles of phages^16,17^ and plasmids^18,19^ in the human gut microbiome, expanding our knowledge beyond the prokaryotic component of communities. These discoveries underscore the importance of investigating the phageome and plasmidome in the skin microbiome, which to date remains largely unexplored.

There have been recent advances in virus and plasmid (mobilome) classification, but distinguishing them from either prokaryotic or eukaryotic host genomes remains challenging^20,21^. Horizontal gene transfer increases the genetic variation of mobile elements, and their frequent integration into diverse host chromosomes further complicates classification, often resulting in low precision from most classifiers^20,22^. Additionally, while bacteria dominate the skin microbiome, microbial eukaryotes, such as fungi, are also essential components of the community, fungal-bacterial interactions significantly influence skin diseases^23^, and human viruses play a pivotal role in modulating the host immune system^24,25^. This multi-domain composition makes the skin microbiome particularly vulnerable to genome binning errors, especially due to fragmented assembly by short-read metagenomics, as the mixture of different domains of life increases the likelihood of contig misassignment across genome bins^26^.

In contrast to the gut microbiome, there have been few large-scale multi-study surveys of skin microbiome diversity^27,28^, and those have been mostly focused on generating catalogues of prokaryotic genomes rather than the determining the factors that drive variation in that diversity. If they have considered phages and viruses, it is without the consideration of potential false positives outlined above. This motivates the careful construction of a multi-domain genome catalogue from skin both to provide an improved reference data set but also by correlating phages with their host strains or other contextual data to better elucidate the role of phages and plasmids in structuring the skin microbiome.

Combining this multi-domain genome catalogue with human genome imputation represents a critical first step toward integrating human genetics and multi-domain microbial interactions, offering new insights into the skin microbiome derived from a single skin swab. It enables us to gain a more complete understanding of the drivers of the skin microbiome: determining the relative importance of human genetics, geography, interactions between microbial kingdoms, and the presence of human viruses.

## Results

### Expanded skin microbial genome collection set from global metagenomics

To generate a comprehensive multi-domain collection of high-quality skin microbiome MAGs we assembled and binned (see Methods) a large collection of publicly available skin metagenomes encompassing 2,264 samples totaling 5.81Tb from five countries (Fig. 1A)^4,5,8,29–31^. Metagenome assembly and genome binning, using CONCOCT, MaxBin2, and MetaBAT2, was conducted separately on single and pooled samples (grouped by the same individual and sampling locations) and integrated through additional refinement steps (see Methods).

**Figure 1.**
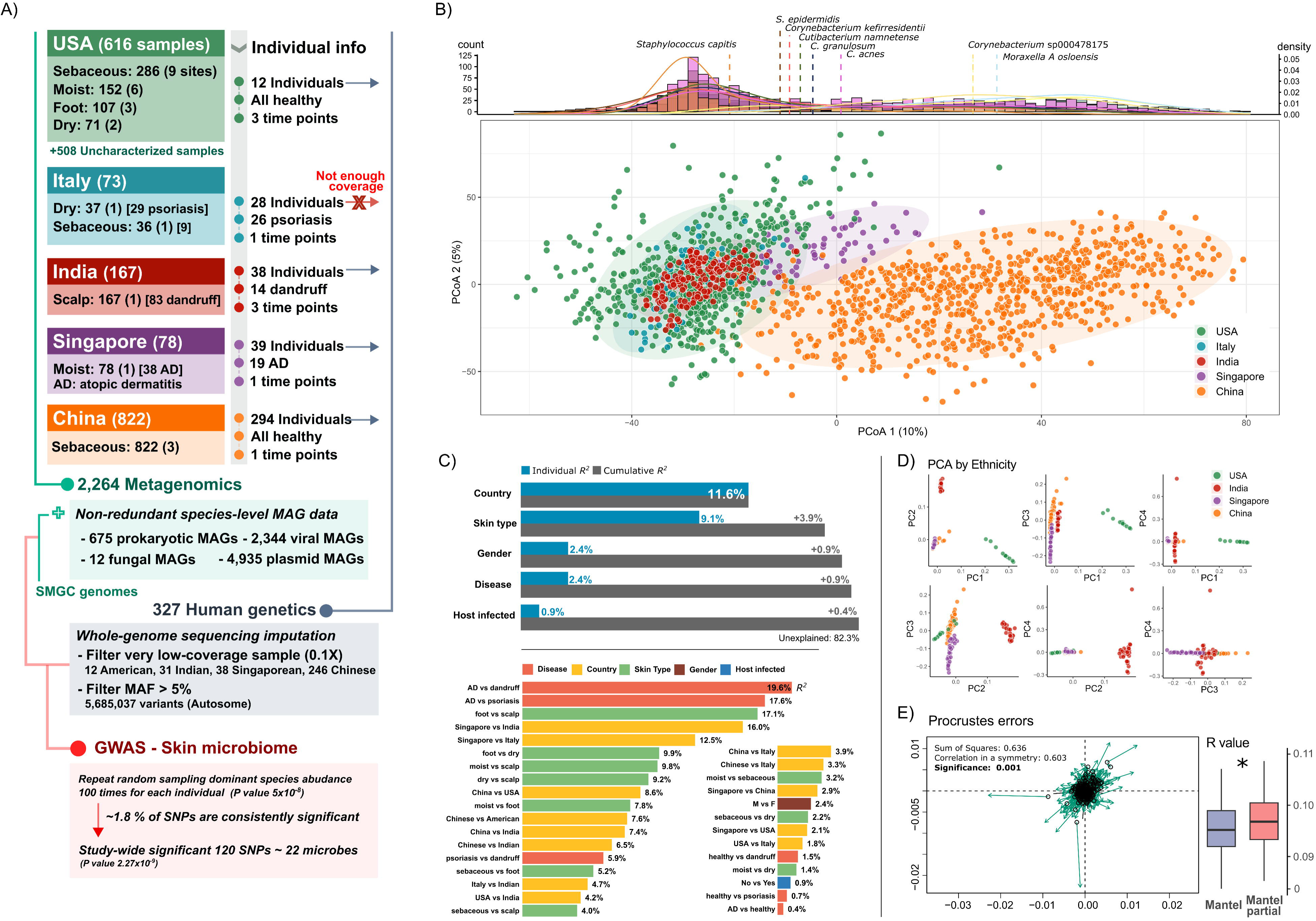
Global skin microbiome analysis strategy and characterization of skin microbial composition across diverse environments. **A)** The workflow illustrating the construction of a multi-domain genome collection and human genetic imputation for GWAS-skin microbiome analyses. The genome catalog was derived from 2,264 publicly available metagenomic datasets, with further analyses performed on 1,756 samples possessing available metadata. Metagenomic data from the same individuals were integrated to impute human genetic information for 327 participants. **B)** Principal coordinate analysis (PCoA) of skin microbiome composition based on Robust Aitchison distances, representing variability across countries. The upper panel displays a density bar plot of microbial species significantly correlated with PC1 values (x-axis), identified by MaAsLin2 analysis^1^. **C)** Individual and cumulative R² values indicating the proportion of microbiome variation explained by environmental factors, including country, skin type, gender, disease status, and host-virus presence. The lower panel presents pairwise comparisons of variables within these environmental factors. **D)** Principal component analysis (PCA) plots depicting human genetic population structure from the four studies using top four principal components (PC1–PC4). **E)** Comparison between skin microbial composition and human genetic population structure. Procrustes analysis was performed on a subset of randomly paired microbial and human genetic data to visualize structural congruence. Mantel and partial Mantel tests, which statistically assess correlations between microbial composition and human genetic structure (*P* value < 0.001), were conducted with iterative subsampling (100 repetitions) of microbiome samples paired with corresponding genetic data. Partial Mantel tests, adjusted for geographic location, showed significantly higher R values compared to Mantel tests (Wilcoxon test, *P* value = 0.024).

The importance of the mobilome in structuring microbial communities is increasingly understood. To reflect this we devised a comprehensive strategy employing multiple classifiers (viralVerify, Virsorter2, plasmidVerify, and geNomad) to recover genomes of previously unknown human viruses, phages, and plasmids rather than relying on a single method (see Methods). Crucially we also filtered for false positives by aligning the predictions of these programs against the NCBI NT database and annotating with 80% nucleotide identity and 40% query length threshold. We applied the classifiers to all contigs > 2kb after clustering at 99% ANI. Of our initial 38,274 predicted viral contigs and 15,687 plasmids, we could annotate 41.64% and 56.08% respectively. Of these annotated contigs, 25.24% of viral contigs and 4.3% of plasmid contigs were found to in fact derive from eukaryotic genome sequences either fungi or metazoans.

Applying this strategy to an existing multi-kingdom genome catalog, the Skin Microbial Genome Collection (SMGC) of Kashaf et al.^28^, revealed similar issues with false positives. Of the 6,935 sequences in SMGC, 35% (2,459 sequences) could be annotated to the NCBI NT database, using the above cut-offs, with 6.1% of these hits (150 sequences) deriving from eukaryotic genomes, primarily fungal and metazoans including human sequences. These false positives included twelve jumbo phages (i.e. 200 kb), which aligned to *Malassezia globosa* CBS7966 and *Malassezia restricta* CBS7877, with average alignment identities of 98.6% and 99.7%, respectively (Supplementary Fig. 1). While the viral genome quality of these jumbo phages was rated as medium by CheckV, we could not find any additional evidence to support these sequences as bona fide viruses or phages based on protein cluster network analysis and gene annotation (Supplementary Fig. 1). Note that these twelve jumbo phages did not include the twenty novel jumbo phages reported in Kashaf et al.^28^.

We integrated our multi-domain genome collection into the SMGC by dereplicating at the species level, but selecting the high-quality option when doing so (see Methods), to create an expanded Skin Microbial Genome Collection (eSMGC). This eSMGC contains 675 prokaryotic MAGs (proMAGs), 12 fungal MAGs (eukMAGs), 2,344 viral MAGs (vMAGs), and 4,935 plasmid MAGs (pMAGs). This represents an 8.3% increase in prokaryotic species number over the SMGC and a substantial improvement in quality, 17.78% of the SMGC proMAGs were replaced, resulting in a significant increase in genome completeness and continuity (paired *t*-test, *P* value < 0.005, average completeness 93.56% and contamination 0.86%, Supplementary Fig. 2). While no novel fungal species genomes were recovered, eSMGC improved eukaryotic genome quality by replacing 50% of the original eukMAGs, resulting in a 1.42% increase in average completeness, a 0.25% decrease in contamination, and a 20.86 kb increase in N50. Similarly, eSMGC discarded low-quality vMAGs and expanded the catalog with 23.91% novel viral genomes, replacing 39.33% of the original SMGC’s viral genomes. To assess the novelty of our virus and plasmid genomes, we compared them to the JGI VR and PR genome databases. At the OTU level (95% identity), eSMGC identified novel 29.86% viruses and plasmids compared to JGI’s most recently updated databases (Supplementary Fig. 2). Finally, we also uncovered previously unidentified human viruses.

### The skin microbiome varies significantly in composition between countries and with skin type

The skin microbiome varies considerably, influenced both by micro-environmental factors, such as physiological skin types^4^, and macro-environmental factors, such as geographical location^6^. To identify the key drivers shaping the skin microbiome, we examined five distinct skin types—sebaceous, moist, dry, scalp, and foot—alongside broader factors, including country of origin, gender, and skin conditions (atopic dermatitis, psoriasis, and dandruff). Additionally, given the known relationship between the skin microbiome and immune status^32,33^, we assessed the impact of host viral infection, classifying each sample as ‘host infected’ if a human virus was detected. Our global study revealed strong country-specific signals in skin microbial composition, which we visualised with a principle coordinates plot (Fig. 1B). The first principle direction (PC1), which correlates with *Moraxella osloensis* and *Corynebacterium* sp., separates Chinese individuals which are enriched for these taxa from the other countries (Fig. 1B, top panel). There is some evidence of clustering, supporting the ‘cutotype’ concept, but the clusters are strongly associated with country. In order to quantify the importance of the different factors that might shape the skin microbiome, we performed multivariate ANOVA of community composition against each factor separately, country emerged as the single most influential variable explaining 11.6% of variation, followed by skin type, gender, disease status, and host-virus presence. We then constructed a combined model by adding factors one at a time starting with country, in order of decreasing variance explained, keeping a variable if it significantly improved the overall fit. The combined model explained a total of 17.7% of the variation (Fig. 1C): with country responsible for 11.6%, skin type 3.9%, while gender, disease status, and host viral presence each added marginal (<1%) explanatory power. If we reorder the variables adding skin type first, its variance explained increases to 9.1% and adding country explains an additional 6.4% indicating that these two variables are correlated. This may reflect the unbalanced structure of the data set, for example scalp and foot samples were only available for the India and the USA, respectively. We therefore restricted the analysis to the dry, moist, and sebaceous skin types which were present for all countries (Supplementary Fig. 3). In this case, country explained considerably more variance (R² = 10.32) than skin type (R² = 4.33). Even when country was added after skin type in this model it still contributed a substantially greater effect (R² = 7.74). Overall, we conclude tentatively, given the limitations of combining studies, that country of origin may be a more important overall driver of skin microbiome composition than differences in skin physiology with type.

### Host genetics impacts skin microbiome composition

This significant impact of country on the skin microbiome could be due to multiple factors, including environment, geography and host genotype. We observed that the two countries mostly comprising individuals of Chinese ethnicity (Singapore and China: Fig. 1B) did appear distinct, suggesting genotype may be important. To fully resolve this association we used genome imputation, which was possible for 327 individuals, to perform host SNP genotyping. Analysis of human SNP differences confirmed that the Singaporean samples were of Chinese ethnicity, as indicated in previous studies^31^, and principal component analysis (PCA) revealed clear differences between ethnicities (Fig. 1D).

We selected 1,541 microbiome samples from 327 individuals for whom host SNP data was simultaneously available. To minimize study-to-study variability in skin sampling sites across countries, we restricted the USA samples—the most extensively sampled—to those anatomical sites available for the other countries (alar crease, cheek, glabella, and antecubital crease). This resulted in a total of 1,047 samples used for subsequent analyses. Due to multiple sites and timepoints we had 3.2 samples per individual on average, to ensure statistical independence we stratified by host, for each host randomly subsampling once from their microbiome samples - repeating this one hundred times.

To quantify the strength of the overall relationship between host genotype and skin microbiome structure, we performed a Mantel test, which demonstrated a significant association between host genetics and skin microbiome composition (Mantel test 100 iterations, an average of R = 0.095, *P* value = 0.001: boxplot inset in Fig. 1E). Even when accounting for country of origin, using a partial Mantel test we found that this significant association remained, with slightly increased magnitude (partial-Mantel test 100 iteration: an average of R = 0.097, *P* value = 0.001: boxplot inset in Fig. 1E).

We performed Genome-wide association analysis (GWAS) between human autosomal variants and the 137 microbiome features present in more than 20% of the 327 individuals (see Methods). The overall GWAS results showed minimal inflation of false-positive associations (median genomic inflation factor λ = 1.02, Supplementary Fig. 4A). Of 67,016 total associations reaching nominal genome-wide significance (*P* value < 5×10^-8^) across these microbiome traits, only 1.8% (1,216 associations) consistently remained significant in more than 90% of iterative association analyses (Supplementary Fig. 4B). These consistently significant SNPs were associated with 22 of the 137 microbiome species, including 5 prokaryotes, 2 fungi, 3 phages, and 12 plasmids. To ensure robust associations, we applied a stricter threshold for study-wide significance (median *P* value < 2.27x10^-9^; Bonferroni corrected for 22 species: 5x10^-8^/22), identifying 107 lead SNPs located within human genes strongly linked to microbial abundance (Fig. 2A). Furthermore, 61 of these SNPs were associated with genes expressed in skin tissue or previously reported in skin-associated variants studies, underscoring their potential relevance (Supplementary Table 1). To determine whether these SNPs remained significant when marginalized by the other factors shaping skin composition. We compared two nested mixed-effect models: one including SNP allele information and one excluding it, while accounting for country, skin type, gender, and disease status. The partial R² values revealed that host SNP alleles independently explained part of the microbial abundance variation, confirming the host genotype’s significant influence on the skin microbiome. Although the R² values were modest, reflecting the complexity of genetic-microbial interactions, the consistent significance of SNP alleles underscores the role of host genetics in relation to skin microbiome (Supplementary Fig. 4C).

**Figure 2.**
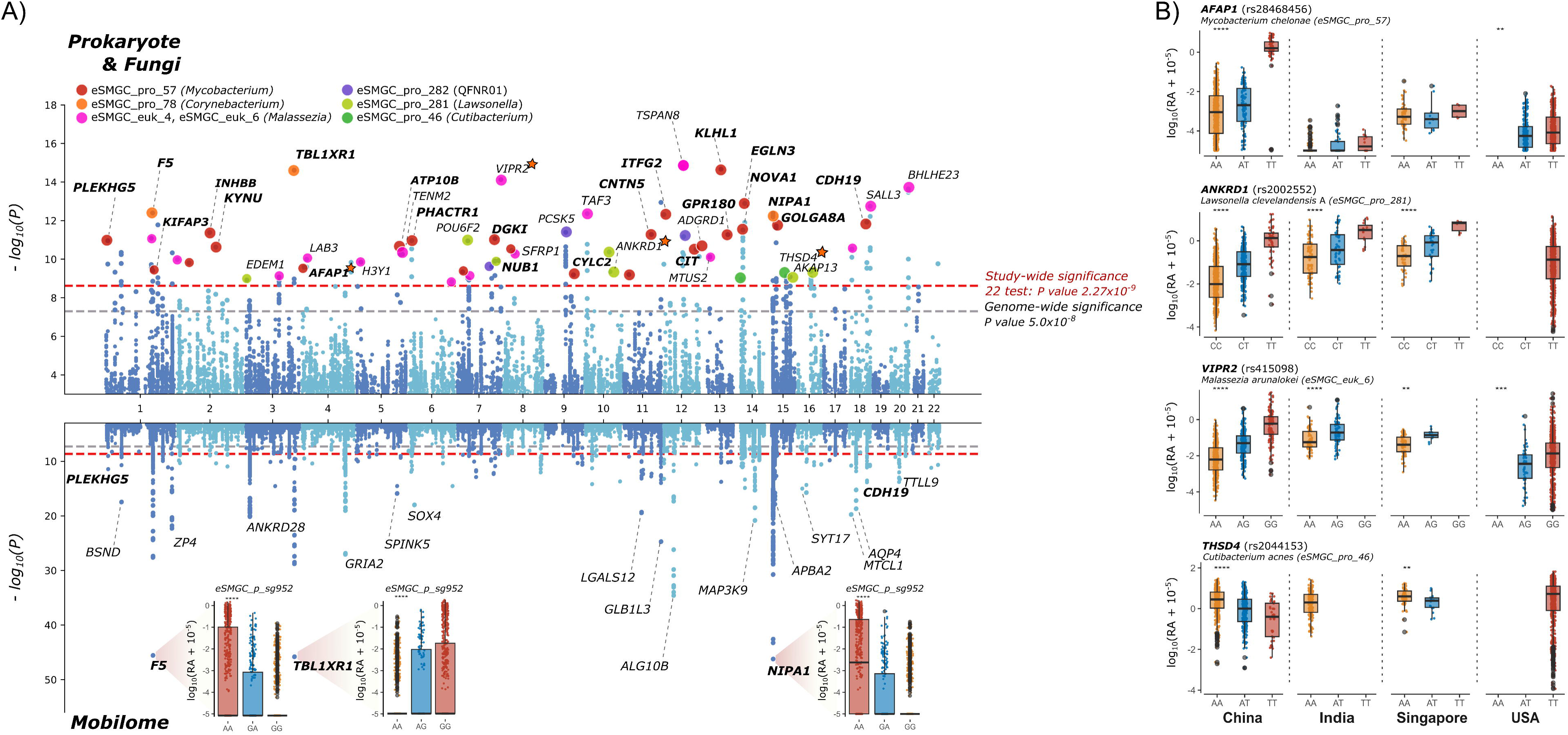
Genome-wide association analysis of skin microbial abundance. **A)** Manhattan plot representing GWAS for the abundance of 22 eSMGC microbial genomes. The upper and lower panels illustrate associations for prokaryotes & fungi (proMAGs, eukMAGs) and the mobilome (pMAGs, vMAGs), respectively. Significance thresholds are indicated by horizontal lines: genome-wide significance (gray line; *P* value = 5 × 10LL) and study-wide significance adjusted for 22 independent MAG tests (red line; *P* value = 2.27 × 10LL). Highlighted points in the upper panel represent lead SNPs, annotated with nearest gene names, associated with major microbial taxa including *Mycobacterium*, *Corynebacterium*, QFNR01 (*Neisseriaceae*), *Lawsonella*, *Cutibacterium*, and *Malassezia*. The inset boxplots in the lower panel display the strongest association observed for an unclassified pMAG (eSMGC_p_sg952) across different host genotypes. Gene names in bold indicate significant SNPs identified in both the upper and lower panels; a star symbol next to the gene name denotes the genes highlighted in panel B. **B)** Examples of consistent microbial association trends across host genotypes from multiple countries, as detailed in Supplementary Table 1.

### SNPs influencing host skin microenvironment correlate with the abundance of prokaryotic species and mobile elements

We observed stronger associations between genomic loci and the mobilome (phages and plasmids) than with host species abundances (Fig. 2A). Mobilomes accounted for 76.2 % of the 214 total significant associations between 107 lead SNPs and 22 microbial traits, with 27% of these demonstrating stronger significance than the most robust associations observed for prokaryotes and fungi (medium *P* value = 2.39x10^-22^; Fig. 2A). The strongest overall association was observed between an unclassified plasmid and the lead variant rs6606827 on the gene NIPA1 (*P* value = 6.19x10^-47^; boxplot inset in Fig. 2A), a gene that may regulate magnesium levels^34,35^ and influence keratinocyte differentiation^36^. Six other unclassified plasmids and one phage sequence also exhibited highly significant associations (median *P* value = 1.13x10^-27^). This variant was also associated with *Corynebacterium* sp. (eSMGC_pro_78; median *P* value = 6.44x10^-13^) remaining consistently significant in exactly 90 of 100 iterative tests, narrowly below our stringent iterative threshold (>90/100 tests). Additional strong associations involving these same mobilomes were found within the F5 locus, where the variant is intragenic, and in the region proximal to TBL1XR1. These loci encode coagulation factor V and transducin beta-like 1 X-linked receptor 1, respectively, and were also associated with *Corynebacterium* sp. (eSMGC_pro_78) abundance.

We identified several loci demonstrating significant associations with the opportunistic pathogen *Mycobacterium chelonae*. These loci encompass genes expressed in melanocytes, fibroblasts, and keratinocytes, which are critical for the integrity and function of skin cell structures, particularly the cytoskeleton, suggesting roles in skin cell regeneration and development. Specifically, CDH19, located near one of the associated SNPs, encodes a type II cadherin that influences cell-cell adhesion, which is crucial for maintaining the skin’s structural integrity. PLEKHG5, featuring a pleckstrin homology domain, actively regulates RhoA to control cytoskeletal dynamics and adhesion in keratinocytes^37^. Additional associations were noted with genes encoding structural proteins such as actin and kinesin. AFAP1, responsible for actin filament-associated protein 1, and PHACTR1, which regulates the disorganization of F-actin, are crucial for maintaining cellular architecture. Following this, KIFAP3, a kinesin II subunit, is implicated in regulating epidermal ciliogenesis. Furthermore, EGLN3, located near the associated variant, encodes a member of the EGLN family of prolyl hydroxylases, which regulate hypoxia-inducible factor 1α activity, thus modulating cellular responses to hypoxia— an essential mechanism for dermal papilla cell proliferation and adaptation under hypoxic conditions^38^. Moreover, *Lawsonella clevelandensis*, a member of the Mycobacteriales order, showed an association with a locus near ANKRD1, which regulates interactions between fibroblasts and the collagen matrix, thereby impacting cutaneous morphology^39^. These results suggest that human genetic variations associated with skin cell structure and morphology may influence the presence of opportunistic pathogenic microbes that are known to infect human soft tissues^40,41^.

In addition, several associations between inflammatory response and the pathogenesis of skin diseases were identified. TLR2 and MAP2K3, involved in skin diseases and inflammations induced by pathogen recognition or stimuli^42,43^, were associated with three mobilome abundances. Additionally, a SNP near KYNU, which encodes kynureninase involved in tryptophan metabolism—a related inflammatory factor in hidradenitis suppurativa^44^ and psoriasis^45^—was associated with *M*. *chelonae*. In parallel, the locus proximal to SFRP1, which downregulates the Wnt signaling pathway suggested to be involved in psoriatic skin^46^, was associated with *Malassezia restricta*. TSPAN8 (tetraspanin 8), located near the associated SNP and recognized as a marker in various human carcinomas^47^ and particularly noted in cutaneous melanoma research^48^, showed a significant association with the abundance of *Malassezia arunalokei*. The abundance of *C. acnes* was significantly associated with the THSD4 locus, previously proposed *in silico* as a potential target involved in Ebola virus infection^49^. Although speculative, this finding supports the hypothesis that *C. acnes* may influence skin immune responses and contribute to host protection by modulating THSD4 expression. Overall, many of these genetic associations with skin microbial abundances were independently replicated in studies from multiple countries, emphasizing the broad relevance of host-microbiome interactions (Fig. 2B; Supplementary Table 1).

### Strain variation in the skin microbiome across countries

We have shown that the overall microbiome composition varies across countries, and that part of this variation is explained by host genotype. We next investigated whether intra-specific microbial variation is also influenced by geography. We focused on abundant species in the eSMGC and determined SNVs using strictly mapped reads (>95% alignment identity) to avoid potential errors caused by mismapping. We identified a total of 4.04 million SNVs in dominant eSMGCs, including 5 eukMAGs, 62 proMAGs, 42 pMAGs, and 7 vMAGs, across 796 samples meeting over 10-fold coverage with >70% genome breadth. These SNVs, supported by considerable read coverage for each sample, allow us to explore the variation landscape and compare skin microbial species across countries (Fig. 3).

**Figure 3.**
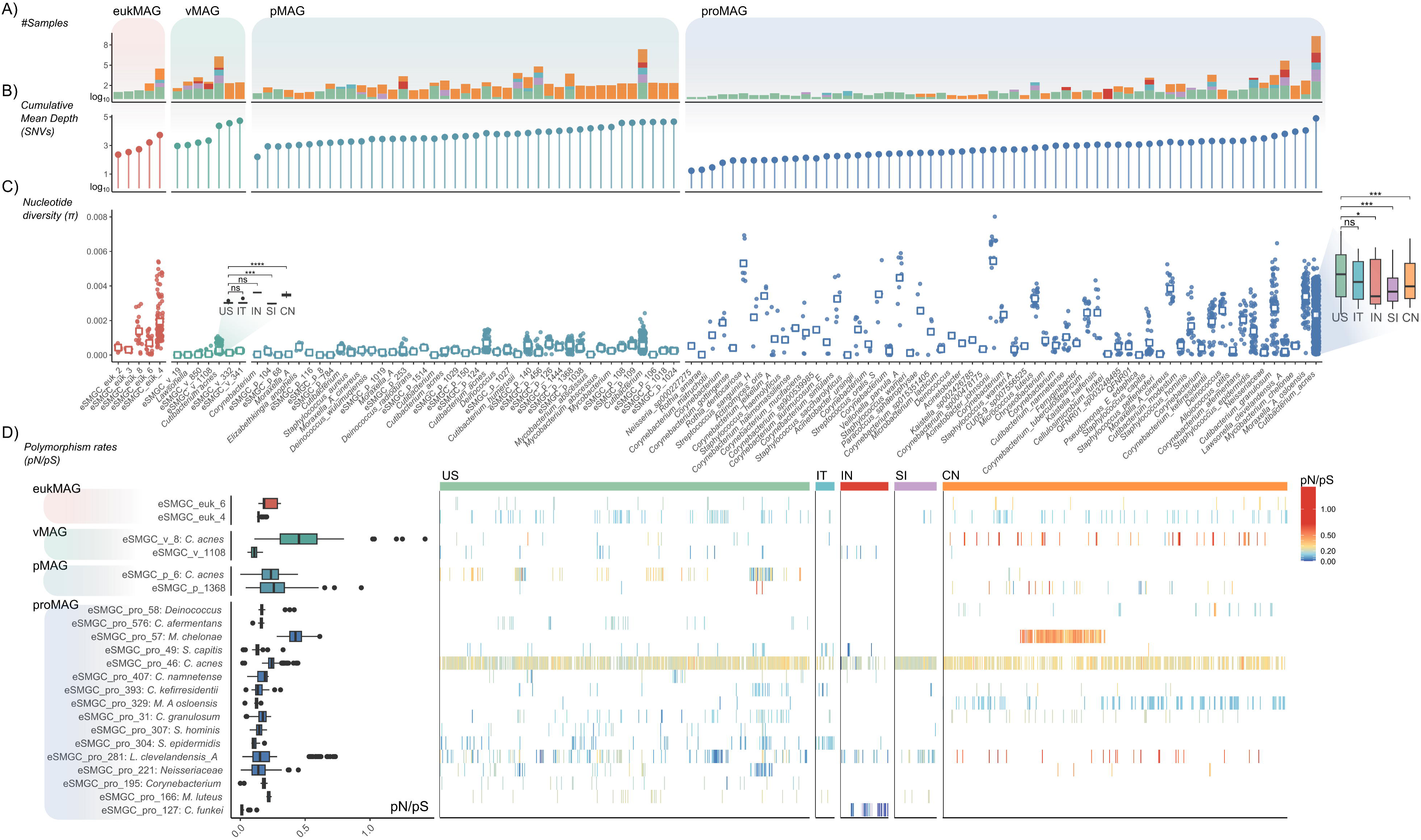
Genomic variation landscape of 116 skin microbial genomes. Genomic variation statistics for 116 skin microbial genomes categorized into four microbial domains: eukMAGs, proMAGs, pMAGs, and vMAGs. Genomic variations were calculated at each genomic position with coverage greater than 10× and genome-wide coverage above 70%. The y-values on each panel indicate: **A)** the number of samples analysed, **B)** cumulative mean sequencing depth, and **C)** nucleotide diversity for each microbial genome exceeding analytical thresholds. Axes in panels **A)** and **B)** were log-transformed (with a pseudocount of 1) for visualization purposes. **D)** Distribution of polymorphism rates (pN/pS) in individual genes for predominant skin microorganisms, showing overall distributions (left panel) and average values per sample across multiple countries (right panel). US – United States; IT – Italy; IN – India; SI – Singapore; CN – China.

Among the most abundant eSMGCs, those deriving from the genera *Cutibacterium*, *Moraxella*, *Mycobacterium*, *Lawsonella*, and their mobilomes, as well as *Malassezia*, had the highest cumulative and mean depth mapped reads for SNV detection (Fig. 3A & B). However, many of these species were only found in a single country preventing cross-country comparisons of their genomic variation (Fig. 3A). These comparisons were possible for *Cutibacterium* and its mobilomes, along with *Lawsonella*, *Corynebacterium*, *Staphylococcus*, and *Malassezia*. For these we calculated microdiversity (π) and polymorphism rates (pN/pS) to determine how genetic diversity and evolutionary pressure vary across species: *M. chelonae* and *L. clevelandensis* had lower intra-species diversity compared to other microbial species while displaying high pN/pS ratios in their genes (Fig. 3C & D). These results may indicate positive adaptive selection in these species. Both species shared common genetic features related to lipid metabolism, including sterol transport proteins and secretory lipases, which likely reflect adaptation to skin environments (Supplementary Table 2). Furthermore, in *M. chelonae*, genes associated with metal binding, zinc/manganese transport system substrate binding proteins (K02077), and oxidative stress (F_420_H_2_-dependent quinone reductase)^50^ showed high pN/pS ratios. Similarly, *L. clevelandensis* had high pN/pS in the PPX/GPPA enzyme (K01524), associated with stress response, similar to findings in *M. tuberculosis*^51^. Selection pressure on genes related to environmental responses and metal uptake may suggest that these opportunistic pathogens have a propensity to rapidly adapt to diverse skin microenvironments.

Notably, *Cutibacterium* and its phage and plasmid genomes had significantly higher levels of nucleotide diversity in Western (USA-Italy) compared to Asian (Singapore-China) regions (boxplot inset in Fig. 3C). However, when examining polymorphism rates, *C. acnes* showed similar patterns regardless of the country, while its phages displayed significantly higher rates in China than in the USA. Especially genes within the restriction-modification (RM) system, a well-known phage defense mechanism, demonstrated more rapid adaptations in Chinese samples than in those from the USA, particularly within the *C. acnes* prophage (eSMGC_v_8) region (Supplementary Fig. 5). These results suggest that the *C. acnes* genome and that of its phages have diverged between countries, we explore this in more detail in the next section.

### Country-wide strain divergence of *Cutibacterium acnes*

A total of 628 *C. acnes* MAGs with >70% completeness and <5% contamination were recovered via co- and individual binning. This was the highest number obtained from any species in our collection. This dataset includes 373 MAGs from the USA, 181 from China, 77 from India, 65 from Singapore, and 10 from Italy. We constructed a phylogenetic tree of these MAGs using universal marker genes. The majority of strains from China formed a distinct clade on this tree (Fig. 4A), whilst the USA strains (deriving from just 12 individuals) exhibited the most phylogenetic diversity probably because of the high number of skin locations sampled. Because of this high variation we used the USA as a baseline and compared *C. acnes* variation in each country against it. We also used this phylogenetic tree to control for population structure by identifying lineage-independent country-specific SNPs.

**Figure 4.**
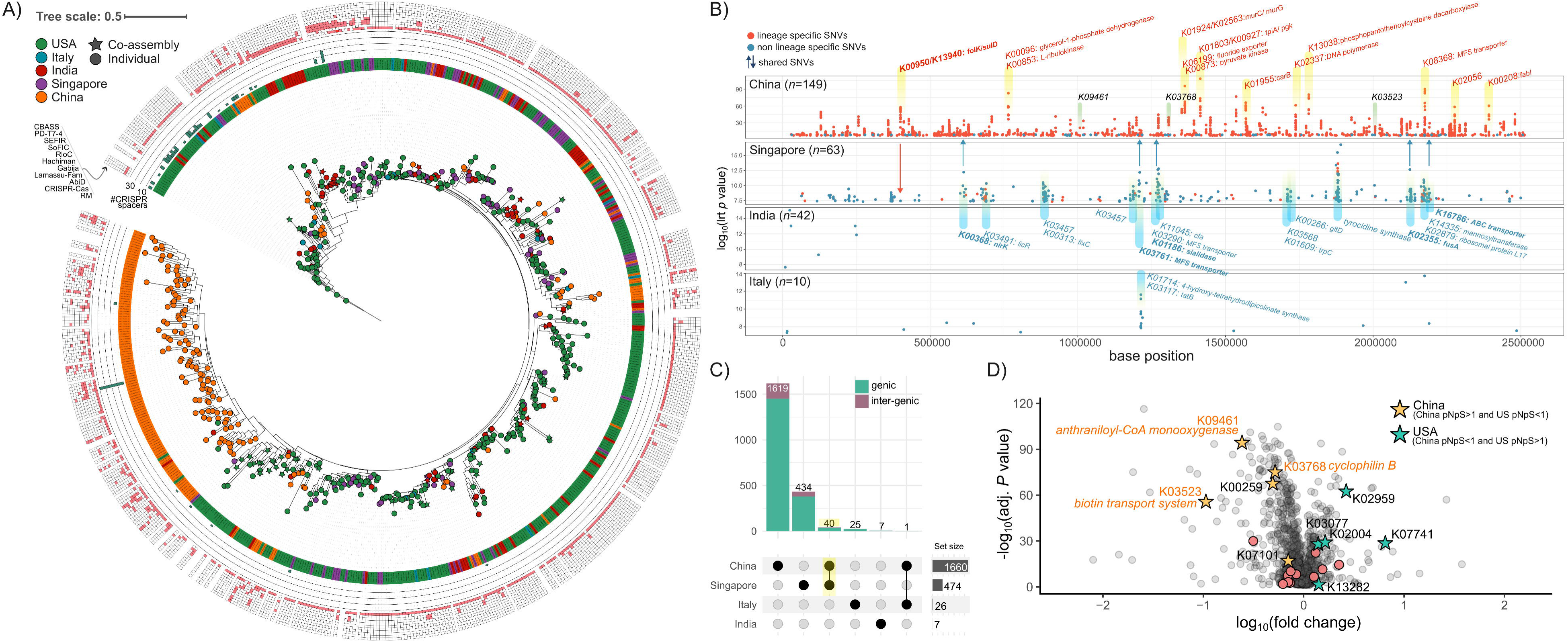
Genomic diversity of *Cutibacterium acnes* strains across multiple countries. **A)** Phylogenetic tree of *C. acnes* MAGs. Nodes and adjacent bars labeled with MAG IDs indicate the country of origin. The next inner ring represents the number of unique CRISPR spacer regions per genome. The outermost ring provides a heatmap indicating the presence of various microbial defense systems within each MAG. **B)** Manhattan plot depicting microbial GWAS comparing *C. acnes* genomes from China, Singapore, India, and Italy to genomes from the USA. All displayed points represent significant SNVs (likelihood ratio test *P* < 5 × 10LL of Pyseer). Lineage-specific SNVs were defined as SNVs significantly associated with both country of origin and phylogenetic structure (*P* < 0.05); all other SNVs were classified as non-lineage-specific. **C)** Upset plot illustrating shared and unique significant SNVs across different countries. **D)** Volcano plot highlighting differential selective pressures between Chinese and USA *C. acnes* genomes, comparing pN/pS ratios of individual genes via Wilcoxon tests (adj. *P* < 0.05). Stars indicate genes with pN/pS ratios >1. Colored KO annotations denote genes that harbor significant SNVs in comparisons between Chinese and USA strains, as presented in panel B.

As expected from the phylogenetic tree, the majority, 96.34% of the SNVs in *C. acnes* significantly associated with China were lineage-specific, whereas most SNVs from Singapore (16.24%), India (0%), and Italy (0%) were not (Fig. 4B & C). Of the total of 2846 country-specific SNVs for *C. acnes* of USA origin, only 638 were lineage-independent, with Singapore having the highest number of lineage-independent SNVs overall, followed by China, India, and Italy. Notably, China and Singapore shared a considerable proportion of lineage-independent SNVs (9.05%) significantly higher than those shared between other countries (India-China 2.28%, Italy-China 1.27%, India-Singapore 0.94%) (Supplementary Fig. 6). This suggests that ethnicity may influence specific genetic variants of *C. acnes* unrelated to lineage, even after accounting for differences in sampling locations (skin characteristics) and national environments.

Furthermore, lineage divergence in *C. acnes* was exclusively observed in Chinese populations (Fig. 4A), where we identified China-specific SNVs compared to the USA (Fig. 4B). The majority of SNVs of Chinese origin were located in genic regions (89.63%), suggesting different roles in environmental adaptation (Fig. 4C). Examining 1500 bp regions flanking the 15 highly significant lead SNVs (lrt *P* value <1x10^-50^), we identified 23 KO-associated SNVs. These were predominantly genes involved in transporter (K08368, K02056), amino acid and nucleotide metabolism (K01955), and lipid metabolism (K00208, K00096) (Fig. 4B). Additionally, we analysed the pN/pS ratios of *C. acnes* genes by SNV between China and the USA to determine which genes were under adaptive selective pressure in this lineage. Significant differences (adjusted *P* value < 0.05) in pN/pS ratios were found in 1137 genes, but the genes that exhibited selective pressure in each country were narrowed down to five in China and USA, respectively (Fig. 4D, Supplementary Table 3). Notably, genes under positive selective pressure in Chinese *C. acnes* included oxidoreductases (K00259, K09461), components of the biotin transport system (K03523), and cephalosporin-C deacetylase (K01060), associated with antibiotic synthesis. In contrast, in the USA, *C. acnes* exhibited selective pressure on ribosomal proteins (K02959, K02954), nucleotide metabolism (K01081), and serine peptidases (K13282), supporting the hypothesis that genetic differences may facilitate adaptation to varying skin macro- and microenvironments (Fig. 4D).

### Species-level diversification of *Cutibacterium acnes* phage by host strain variation

Genetic variants of *C. acnes* were found to adapt to different skin environments, but we also found that this phylogenetic differentiation was strongly intertwined with their phages. Phages and viruses are more individual, site-specific, and environmentally sensitive than other organisms (Supplementary Fig. 7). After building eSMGC, a more definitive collection of skin microbial genomes, the reconstructed 126 *C. acnes* phage genomes were specifically distinguished at the species level in the overall phage phylogeny (Supplementary Fig. 8). In particular, the core genes phylogenetic tree separated them into two clades clearly, with clade-II being a particular differentiation in the Chinese sample (Fig. 5A). We found that clade-I consisted of two subclades (clade-Ia, Ib), with clade-Ia being USA specific and -Ib being found in Chinese and Singaporean samples, corresponding in their abundance (Fig. 5A, middle panel). Furthermore, matching the samples in which these phages were present with the corresponding sample-specific *C. acnes* strains identified by individual binning (Fig. 5A, bottom panel; the pruned phylogenetic tree of Fig. 4A) showed that the phylogeny of *C. acnes* and these phages were highly congruent (ParaFit, *P* value < 0.001). Many of these showed cross-country differences between Chinese and USA samples that did not coexist simultaneously. Notably, the USA samples with eSMGC_v_240 on clade-II were mainly from the same individual, suggesting that phage specificity may reflect not only host lineage but also individual environmental characteristics.

**Figure 5.**
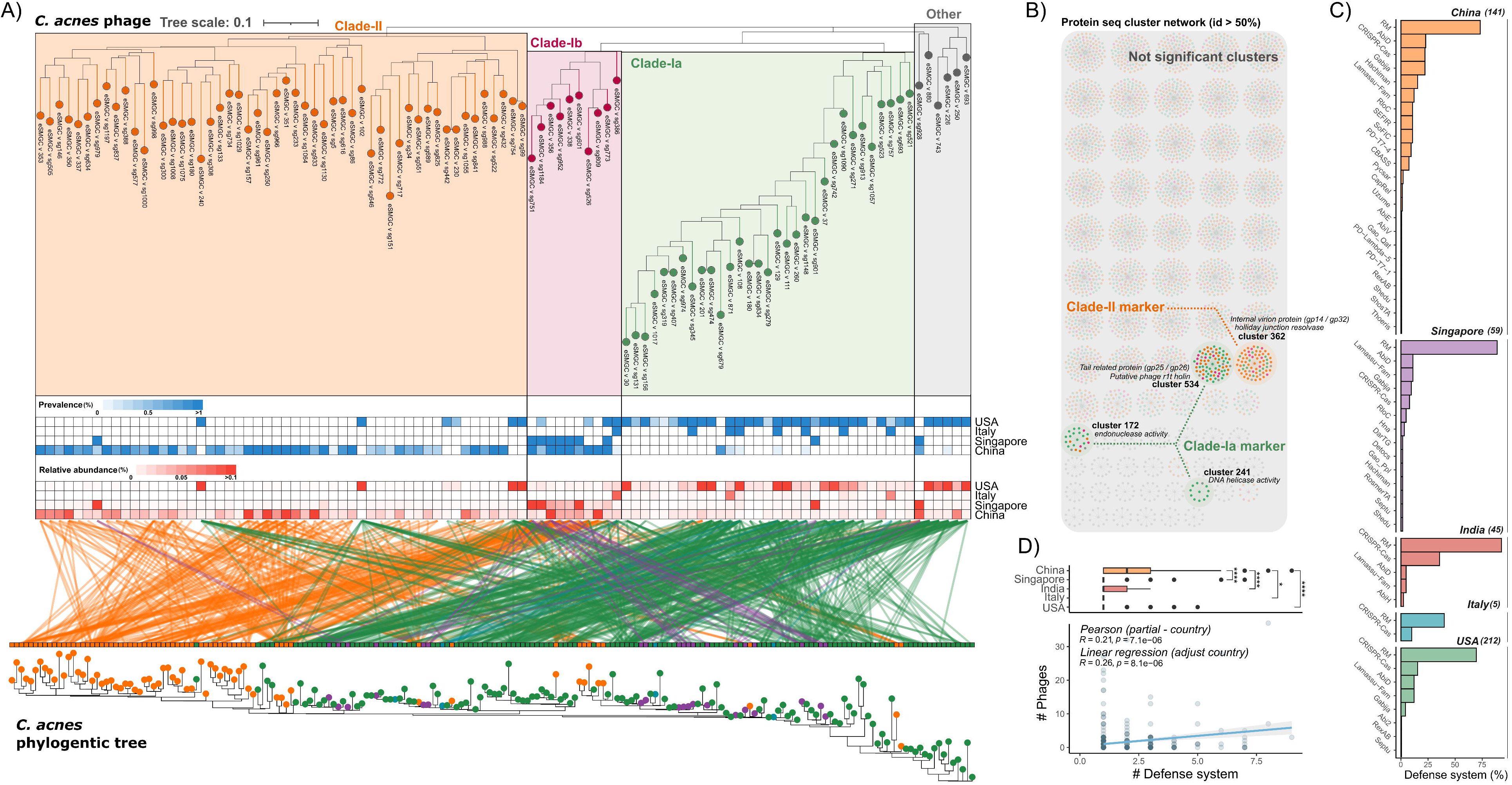
Diversity of *Cutibacterium acnes* phages across host strains and geographic locations. A) Phylogenetic tree of *C. acnes* phages constructed from shared gene content. Middle heatmap panels shows phage prevalence and relative abundance across samples from multiple countries. The bottom panel illustrates associations between specific *C. acnes* strains and their respective phages, reflecting congruent alignments with corresponding phylogenetic trees. **B)** Protein-protein cluster network illustrating the presence or absence of pan-protein clusters in each *C. acnes* phage clade. Specific clade-marker protein clusters are highlighted, as identified by Fisher’s exact test (adj. *P* < 0.05). **C)** Distribution of phage defense systems in *C. acnes* strains originating from each country. **D)** Linear regression analysis evaluating relationships between the number of *C. acnes* phages and defense systems within individual samples. The upper boxplot panel compares the number of defense systems across different countries.

Although *C. acnes* phages generally feature densely located protein-coding sequences with similar genomes^52^, we conducted a pangenome analysis to explore differences in genetic characteristics across lineages. From 126 phage genomes (mean genome size: 29kbp), protein sequences were clustered into 562 nonredundant clusters with at least 50% amino acid identity (Fig. 5B). Notably, four gene clusters (cluster172, cluster241, cluster362, cluster534) showed significant variances between clade-Ia and -II, while no differences were observed between clade-Ib and -II (Fisher’s exact test: adj. *P* value <0.05). Cluster241 (DNA primase) and cluster362 (PA6 phage gene product [Gp] 33) were predominantly found in clade-Ia and -II, respectively. These clusters, being adjacent in the genome, suggest that genetic variations may influence each other’s open reading frames. Cluster534 (Gp26) was absent in some clade-II phages but present in most clade-Ib. Sequence variations between the holin and Gp23 genes, adjacent to cluster534, likely contribute to these genomic differences (Supplementary Fig. 9). Cluster172 (Gp45), located at the terminal end of the phage genome, was inferred to be missing from insufficient metagenomic assembly, indicating limitations in distinguishing phage clades. Although the functions of most PA6 Gp remain unknown, these genetic variations can affect genes like DNA primase and holin, potentially altering phage proliferation strategies against different host strains.

As bacteria possess diverse antiviral defense strategies to prevent phage invasion^53,54^, we sought to understand their association with phages by examining antiviral defense systems in *C. acnes* genomes recovered from individuals. 462 (91.5%) of *C. acnes* genomes contained at least one defense system. The most commonly identified defense system was RM, present in approximately 75% of genomes, followed by CRISPR-Cas, Abi, Lamassu, and Gabija (Fig. 5C). Furthermore, we observed phylogeny-enriched patterns in specific defense strategies: some *C. acnes* genomes contained only CRISPR-Cas, others possessed both CRISPR-Cas and RM, some featured Abi, Lamassu, and Gabija, and the majority contained only RM (outer ring of Fig. 4A). Additionally, the number of defense systems in each *C. acnes* varied greatly, ranging from 1 to 9, and the number of genes varied from 1 to 24. Notably, most *C. acnes* derived from China had a significantly higher number of defense systems than those from other countries (Fig. 5C & D), regardless of their lineage (outer ring of Fig. 4). These genomes also exhibited a wide variety of defense systems, including Hachiman, SEFIR, SoFIC, and CBASS (Fig. 5C), which were not frequently found in genomes from other countries. In line with previous observations that reported *C. acnes* genomes possess an I-E type CRISPR-Cas system^52,55^, we observed that CRISPR systems in *C. acnes* genomes from all countries were of the I-E type; however, only those from China exhibited an I-G type, which has not yet been reported. Next, we matched coexisting phages to each *C. acnes* by individuals to explore whether different defense systems enhanced or protected against phage invasion. We discovered a significant positive correlation between the number of coexisting phages and the complexity of defense systems, after correcting for the country effect (Fig. 5D) (*R* = 0.26, *P* value = 8.1 x 10^-6^) rather than identifying a distinct pattern of phage coexistence based on specific defense systems (Supplementary Fig. 10).

### Host virus presence affects skin microbiome diversity

In Fig. 1C, the presence of host viruses was identified as an additional factor influencing skin microbiome structure, independent of country, gender, or disease state, prompting further exploration of this effect. As the sequencing depth between samples could bias microbial diversity, we selected sufficiently sequenced samples (>10M paired-end reads) and compared microbial diversity in the presence and absence of host viruses. Of the total 1,756 samples, 921 samples from the USA and China met the threshold of 10M paired-end reads, resulting in a balanced distribution of samples with (China: 346, USA: 135) and without (China: 284, USA: 156) host viruses in each country (China: 45.1% vs. 54.9%; USA: 46.4% vs. 53.6%), providing a suitable dataset for comparison.

We determined that 382 (16.3%) of the 2,344 vMAGs represented mammalian viral genomes capable of infecting host cells, predominantly from the *Papillomaviridae* family (*n*=373, 97.64%), followed by *Polyomaviridae* (5), *Poxviridae* (2), *Retroviridae* (1), and unclassified *Sepolyvirales* (1). Compared to the Papillomavirus Episteme (PaVe) database of the HPV genome, eSMGC included 58 previously uncharacterized HPV genomes at vOTU level, comprising 18 novel types and one novel species. In addition, three vMAGs exhibited less than 60% nucleotide identity in the L1 gene and were classified as feline papillomaviruses, indicating their assignment to a distinct genus and non-human host. The *Papillomaviridae* genomes derived from diverse genera including gamma, beta, and alpha papillomaviruses, with a higher prevalence of beta papillomavirus across individuals compared to gamma papillomavirus (Fig. 6A; mean of 6.15 for beta per individual and 1.14 for gamma per individual), despite a lower diversity in beta papillomavirus. Intriguingly, feline papillomavirus was consistently detected in one individual across different sampling times and sites (Supplementary Fig. 11A), despite no reported cases of zoonotic transmission, suggesting continuous human exposure probably as a result of that individual’s particular environment.

**Figure 6.**
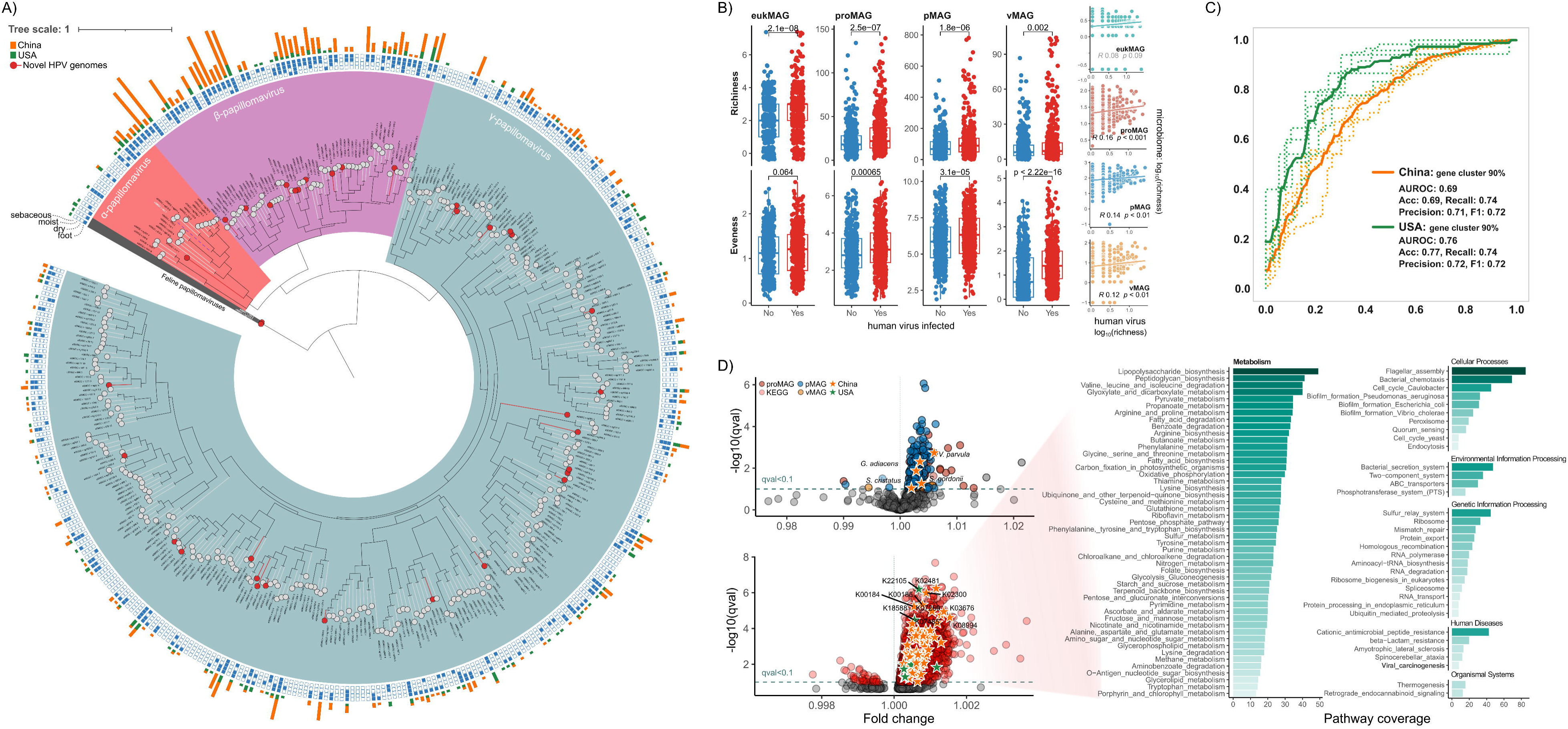
Host-virus genome diversity and impact on skin microbial diversity. **A)** Phylogenetic tree of *Papillomaviridae* genomes collected from eSMGC, including both human and feline papillomavirus. The inner ring indicates the skin types where each virus was detected. The outer ring (bar plot) represents the number of individual participants harboring each papillomavirus genome. **B)** Comparison of skin microbial diversity in relation to host-virus presence (Wilcoxon test). The right panel illustrates the linear relationship between microbial diversity and human viral diversity. Note that vMAG diversity in panel B excludes host-virus diversity. C) Receiver operating characteristic (ROC) curves classifying human viral presence in Chinese and USA samples based on 5-fold cross-validation. Mean ROC curves are shown as bold lines, with corresponding classification statistics summarized in the bottom-right corner. **D)** Volcano plots highlighting marker MAGs and genes differentiating microbial communities based on host-virus presence (adj. *P* < 0.1; by MaAsLin2 analysis^1^). Stars indicate selected marker features selected in XGBoost classifier models for Chinese and USA samples in panel C. The right panel shows enriched metabolic pathways reconstructed from significant genes. 1 Mallick, H. et al. Multivariable association discovery in population-scale meta-omics studies. PLoS computational biology 17, e1009442 (2021).

We also found one sequence identical to the human immunodeficiency virus (HIV) genome, which we assigned as eSMGC_v_851. This was consistently found in samples from the same individual (HV11) across different sampling sites, times, and even sequencing libraries in the Oh et al.’s study^4,5^ (average of 28.8% genome coverage at 0.71 read depth; Supplementary Fig. 11). This finding suggests a potential HIV infection in subject HV11, despite the participants in the Oh et al. studies being healthy volunteers.

The presence of host viruses was associated with a highly significant increase in microbial diversity across all four domains and the richness of host viruses was positively correlated with microbial richness (Fig. 6B). This was not just due to inter-individual differences, within individual sites with host viruses present had higher diversity as confirmed through paired t-tests (Supplementary Fig. 12). To better understand the links between host viruses and microbiome composition and diversity we developed XGBoost machine learning model to predict virus presence from microbiome taxa and functional gene clusters. We built independent models for China and the USA, to account for the major differences in microbiome between these two countries, achieving mean AUCs of 0.65 (±0.01) and 0.74 (±0.02) respectively, indicating significant predictive power (Fig. 6C). For the Chinese and USA sets, the models identified 67 and 129 features, respectively, with 38 in China and 40 in the USA being significant in more than 70% of the iterations. Notably, there was no overlap in features between the two countries. Important features for China included *Streptococcus gordonii*, *Granulicatella adiacens*, *Empedobacter falsenii_A*, *Veillonella parvula_A*, *Serratia grimesii*, and 33 gene clusters, whereas for the USA, *Staphylococcus auricularis*, 12 pMAGs, and 27 gene clusters were significant (Fig. 6D). These features were further validated by Multivariate Association with Linear Models 2 (MaAsLin2) analysis^56^, confirming that most were significant (Fig. 6D).

We annotated the gene clusters to KEGG orthologs (KO) to determine the functions impacted by the presence of host viruses. Reflecting the diversity increase associated with the virus, more significant KOs positively correlated with virus presence (Fig. 6D, 2,928 KOs, adj. *P* < 0.1), than were negatively associated (Fig. 6D, 40 KOs, adj. *P* < 0.1). These enriched pathways included amino acid and pyruvate metabolism, but also modules associated with human innate immunity, particularly pathways involving bacterial cell components, such as flagellar assembly, lipopolysaccharide, and peptidoglycan biosynthesis, critical for detecting invading pathogens via pattern recognition receptors (Fig. 6D)^57–59^. Additionally, the cationic antimicrobial peptide resistance pathway was enriched in these samples, along with biofilm formation pathways, suggesting strategies that microbes employ to evade innate immune responses.

## Discussion

This is the first study to apply ultra-low-coverage human genetic imputation to resolve both host genotype and the microbiome simultaneously from human skin. We expect this to become a common strategy that will be applied in future to other body sites. By performing a multi-domain meta-analysis across multiple countries, we were able to demonstrate that geographic differences in microbiome composition exceeded previously reported levels and were comparable to the differences between physiologically different body sites. Importantly, some of the unexplained variation in skin microbiome structure can now be attributed to host genetics—a small but statistically significant proportion, approximately 1%. While the overall effect size is modest, several microbial taxa exhibit strong associations with host genotype. Our GWAS results identified 22 microbial taxa spanning bacteria (e.g., *Cutibacterium*, *Corynebacterium*, *Mycobacterium*), fungi (*Malassezia*), phages, and plasmids, with robust and novel associations involving 107 human genetic loci.

Notably, several of these associations were replicated across different countries, providing convincing evidence that skin microbial abundance is impacted by human genotype. This is plausible, as host SNPs can influence skin physiological traits^11,60,61^ including earwax type, sebum production, hydration, and hair follicle development, and hence the niche-specific composition of the skin microbiome. We found stronger associations between human SNPs and elements of the mobilome, phages and plasmids, than we did for microbial taxa either prokaryotes or fungi. This is interesting it underscores the importance of MGEs as a means for microbial species to rapidly adapt to host-driven environmental changes, as has been observed in the gut microbiome^19^. The actual mechanisms by which host genetics are shaping the skin microbiome and the colonization of specific microbes and MGEs cannot be resolved by statistical associations alone, and further research will be needed to elucidate these links.

It is interesting to compare these results with what is known for the associations between the better-studied gut microbiome and human genotype. There are relatively few comparable gut microbiome studies with host genetics and metagenomes that span multiple countries. Studies involving single population cohorts have identified substantial numbers of significant SNPs e.g. 567 independent associations from nearly 6,000 paired metagenome and fecal metagenomes^62,63^. Some associations have been confirmed across multiple studies, for instantly that between the *LCT* lactase locus and *Bifidobacterium* albeit that interacts with dairy intake. Recently, it has been shown that some intra-species gut microbiome variation correlates with host genotype notably in *Faecalibacterium prausnitzii* the presence of a N-acetylgalactosamine (GalNAc) utilization gene cluster is associated with a host phenotype that is jointly determined by human ABO^64^ and FUT2 genotypes^65^. The latter controls the fucosylation of mucosal glycans and variants in this gene can result in other gut microbiome changes implying it may have a broad impact on gut niches. We found intra-strain variation in *C. acnes*, but this variation appeared primarily attributable to environmental and ethnic differences between individuals from China and the USA, rather than directly linked to specific host genetic functions as described in gut microbiome examples. After correcting population structure, genetic and structural variations in *C. acnes* were associated with a small number of host SNPs at marginal statistical significance thresholds (Supplementary Fig. 13). These SNPs have not previously been reported in microbiome studies, and their biological significance remains uncertain. Nevertheless, given the significant impact of host genotype on species composition identified here, future studies utilizing large-scale multi-ethnic datasets across countries may clarify these associations and further elucidate genotype-driven skin microbiome variations.

The bacteria-phage relationship emerged as another critical factor in explaining the clear phylogenetic differentiation of *C. acnes* populations between Chinese and other samples. While *C. acnes* exhibited intraspecies-level differences across countries, its associated phages displayed interspecies-level phylogenetic divergence, indicating that phages undergo more rapid genomic evolution compared to their bacterial hosts. This divergence appears to drive *C. acnes* to adopt distinct defense strategies against phages in different populations. Although it remains unclear whether bacterial or phage evolution drives these dynamics, our findings emphasize their mutual influence and suggest that bacteria-phage relationships should not be analysed as strictly comparable taxonomic ranks. Understanding these relationships at the intraspecies level could provide valuable insights for the future of phage therapy, potentially enabling strain-specific modulation tailored to different countries and ethnicities to support a healthy skin microbiome.

The skin functions not only as a physical barrier but also as a vital immunological defense essential for preventing wounds and infections. Eukaryotic viruses transmitted through skin contact can disrupt the host’s innate immune system by infecting epidermal keratinocytes and dermal fibroblasts. Previous studies have shown that healthy individuals typically exhibit a relatively low diversity of eukaryotic viruses in their skin microbiome, whereas individuals with immunodeficiency display markedly increased viral representation and diversity^32^. This suggests a close relationship between skin-associated viruses and the immune system. However, the impact of these viruses on the skin microbiome remains poorly understood, particularly in healthy individuals who may harbor such viruses asymptomatically.

Our study observed a trend of increased microbial diversity in samples containing host viruses. Several factors may explain this finding. First, the majority of viruses identified were low-risk HPVs, which are known to infect keratinocytes and can delay their differentiation^66^. This disruption in the normal skin cell maturation process may alter the skin microenvironment, creating niches that support the growth of a broader range of microbial species, thus increasing overall diversity. Second, we detected a significant enrichment of microbial functional genes associated with antimicrobial peptide resistance, flagellar assembly, and biofilm formation in the presence of eukaryotic viruses. While biofilm formation by certain bacteria like *S. aureus* can reduce microbial diversity by facilitating the dominance of specific species^67^, the overall effect in our samples was an increase in diversity. This suggests that, in the context of eukaryotic virus presence, biofilm formation and flagellar motility may enable a variety of microbes to colonize the altered skin environment more effectively^68^. Furthermore, the presence of HPV may activate the local immune response^69–71^, such as antimicrobial peptides like LL-37 and β-defensins, which can selectively inhibit some microbes while allowing resistant species to thrive^72^. These changes could alter microbial composition and potentially increase microbial diversity by creating niches for less dominant species.

In addition to scientific insights, our analyses revealed some important methodological considerations when performing multi-domain analyses of low microbial biomass samples such as those from skin. We demonstrated the importance of carefully validating phage predictions, identifying multiple false positives in the existing Skin Microbial Genome Collection (SMGC)^28^ and producing our own better curated expanded SMGC (eSMGC) which we hope will be a useful tool for other researchers. Furthermore, our observation of the human HIV genome in multiple samples from the same individual in the Oh et al. study demonstrate the potential usefulness of skin metagenomes for detecting host infections.

To advance research into skin microbiome-host associations, rigorous standardization of workflows is crucial. Implementing host DNA depletion methods to balance microbial and host DNA recovery—rather than sequencing them separately—could enhance efficiency and provide deeper insights into host-microbe interactions. Additionally, the development and curation of a comprehensive, multi-domain skin microbiome database, combined with the integration of recently published large-scale metagenomic studies^73,74^, will be invaluable for strengthening our findings and uncovering the complex interactions between host genetics, environmental factors, and microbial communities. These advancements will significantly enhance our understanding of the factors shaping skin health and disease, paving the way for microbiome-based therapeutic applications.

## Methods

### Public metagenomic data collection

We collected 2,264 public skin metagenome data generated from six different studies across five countries: the United States (1,124 samples)^1,2^, Italy (73)^3^, India (167)^4^, Singapore (78)^5^, and China (822)^6^. These datasets were derived from 403 diverse volunteers, with the breakdown per country as follows: the United States (12 volunteers), Italy (21), India (38), Singapore (39), and China (293). The distribution of samples for each study is detailed below:

#### United States^1,2^

Metagenomic data were collected at three different time points from 18 sampling sites, including the alar crease (Al), cheek (Ck), glabella (Gb), external auditory canal (Ea), retroauricular crease (Ra), occiput (Oc), back (Ba), manubrium (Mb), nare (Na), antecubital crease (Ac), interdigital web (Id), popliteal fossa (Pc), inguinal crease (Ic), toe webspace (Tw), plantar heel (Ph), toenail (Tn), volar forearm (Vf), and hypothenar palm (Hp).

#### Italy^3^

Metagenomic data were collected from two sampling sites, Oc and Ra, corresponding to dry and sebaceous skin types, respectively, at a single time point. In Italy, each individual provided paired samples—one from a skin site with psoriasis and one from a healthy skin site.

#### India^4^

Metagenomic data were collected from the scalp (Sc) at three different time points. Samples were obtained from individuals with or without dandruff.

#### Singapore^5^

Metagenomic data were collected from the Ac at a single time point. Samples were obtained from individuals with or without atopic dermatitis (NCBI accession number: SRR1950743 ∼ SRR1950742).

#### China^6^

Metagenomic data were collected from healthy individuals at three skin sites (Al, Ch, and Gb) all corresponding to sebaceous skin type, at a single time point (CNGBdb accession number: CNP0000635).

Of these samples, metadata (including personal ID, country, sampling time point, sampling site, and disease status) were available for 1,781 samples (Supplementary Table 4), facilitating further association analyses. For analysis purposes, all sampling sites were categorized into five skin types: dry (Hp, Vf), moist (Ax, Ac, Ic, Id, N, Pc), sebaceous (AI, Ba, Ch, Ea, Fh, Gb, Mb, Oc, Ra), scalp (Sc), and foot (Ph, Tn, Tw).

### Data preprocessing and genome assembly

Metagenomic sequences were trimmed using FaQCs^7^ (v.2.09) with the default option. Subsequent removal of human sequences (hg38) was performed using NCBI BMTagger^8^, as recommended by the HMP project. Sequences from the Li et al. study^6^, already pre-processed (quality-trimmed and with human sequences removed), were used directly for metagenomic assembly. Quality filtered reads were assembled using metaSPAdes^9^ (v.3.15) with -k 21, 33, 55, 77, 99, 121. For best optimization, we constrained the maximum value of the k-mer option based on the median length of each sample (k 99 or k 121). Additionally, we incorporated pre-calculated metagenomic assemblies from both 2014 of Oh et al. and Chng et al. studies^1,5^, available at http://segatalab.cibio.unitn.it/data/Pasolli_et_al.html. Co-assemblies were performed on samples sharing biogeographical characteristics from at least three samples to maximize the recovery of low-abundance microbial sequences. Specifically, samples from the same sampling site in the Oh et al. study, those belonging to the same skin type per individual in the Oh et al. study^2^, and scalp samples per individual in the Saxena et al. study^4^ were pooled for co-assembly.

### Genome binning and curation

To calculate the abundance of each assembled contig, quality-filtered reads were mapped using Bowtie2^10^ (v2.3.4.3) with default settings, and the resulting alignments were sorted with Samtools^11^ (v1.8). Genome binning was performed using three different tools—CONCOCT^12^ (v1), MaxBin2^13^ (v2.2.4), and MetaBAT2^14^ (v2.2.15)—following the default procedures described on their respective websites (https://concoct.readthedocs.io, http://downloads.jbei.org/data/microbial_communities/MaxBin/README.txt, and https://bitbucket.org/berkeleylab/metabat). Prokaryotic and eukaryotic genome bins were further refined using ACR^15^ (v0.2) with the options --target both (for both prokaryotes and eukaryotes) and -j Y (utilizing JGI coverage files), and subsequently filtered with GUNC^16^ (v1.02, default options) to remove chimeric bins. For virus and plasmid classification, viralVerify^17^ (v.1.1), VirSorter2^18^ (v2.2.4), and plasmidVerify^19^ were first applied separately to individual and co-assembled contigs, considering only contigs >2 kb. Redundant sequences (> 99 identity) were then removed using CD-HIT-EST^20^ (v4.8.1; -c 0.99 -n 10). The resulting non-redundant viral and plasmid contigs were re-classified using geNomad^21^ (v1.1.0; default options) and further filtered to retain only those with scores above 0.8 for both virus and plasmid categories. To correct for potential misclassification, these contigs were blasted against the NCBI NT database (BLASTn^22^ v2.9.0+; -evalue 1e-10 -perc_identity 80 -outfmt ‘6 std qlen slen’). Contigs with aligned coverage exceeding 40% and alignment identity greater than 80% across all significant BLASTn^22^ hits were assigned taxonomic ranks based on NT sequences. If any alignment classified a contig as Eukaryota, it was excluded from further analysis. Similarly, if any alignment classified a contig as Virus but the contig had a geNomad^21^ score below 0.8, it was recovered for further analysis as a mobilome. For viral contigs, CheckV^23^ (v0.7.0, default options) was used to assess quality, and only contigs with medium, high, or complete quality were retained. Additional proviral sequences were recovered via CheckV^23^ and labeled with “_multicopy” if the repeated sequence length exceeded 1 kb and the product of the repeated sequence length and copy number exceeded 60% of the total contig length. Viral contigs of SMGC^24^ were re-classified as virus and plasmid using geNomad^21^, following the same correction steps, for incorporation into the subsequent eSMGC catalog.

### Expanded Skin Microbial Genome Collection (eSMGC) construction

For representative proMAG selection, we used dRep^25^ (v3.2.0; -comp 50 -cont 10 -sa 95) with the ‘dereplicate’ command for medium- and high-quality MAGs at ≥95% ANI. Prokaryotic MAGs from SMGC were incorporated and further dereplicated using the same parameters. Taxonomic classification was performed using GTDB-Tk^26^ (v2.0.0) with the classify_wf module, and a phylogenetic tree was generated using PhyloPhlan3^27^ (v3.0.64; --diversity high --fast).

For representative eukMAG selection, dRep^25^ (v3.2.0; -pa 0.9 -sa 0.95 -nc 0.30) ‘compare’ was used based on genome quality scores calculated by EukCC^28^ (v.0.1.5; database: eukcc_db_20191023_1). Representative eukMAGs were then selected using a genome quality score defined as: completeness – 5×contamination + 0.5×log(N50). Eukaryotic MAGs from SMGC^24^ were incorporated and processed similarly.

To select representative mobilome clusters, vMAGs and pMAGs were clustered at the population level (≥95% ANI across ≥85% aligned fraction)^29^ via all-vs-all pairwise BLASTn comparisons. Re-classified viral and plasmid contigs from SMGC^24^ were then incorporated into the vMAG and pMAG clusters using the same thresholds. Final mobilome clusters were designated as eSMGC_v_XX for viral clusters or eSMGC_p_XX for plasmid clusters, with singletons prefixed with “sg” Taxonomic classification was conducted using geNomad^21^, and a draft phylogenetic tree was generated using ViPTreeGen^30^ (v1.1.2). To verify the novelty of the eSMGC mobilome, we compared clusters at the population level (95% identity) with the JGI IMG VR4 and PR genome databases^31,32^ using BLASTn. Additionally, genus-level host taxon predictions were processed with iPHoP^33^ (v1.2.0, default options) using an updated iPHoP^33^genome database that incorporated eSMGC’s proMAGs.

Subsequent gene prediction and annotation of eSMGC MAGs were performed to capture the domain-specific characteristics. Specifically, Prodigal^34^ was employed for proMAGs and pMAGs, Prodigal-gv^21^ for vMAGs, and GeneMark-ES^35^ for eukMAGs. Predicted genes were then functionally annotated using EggNOG mapper^36^ (v2.1.12), which provided comprehensive information on gene functions and orthology assignments within the eSMGC.

### Microbiome profiling

Metagenomic reads were aligned against the eSMGC MAG database using Bowtie2^10^ (v2.3.4.3, --very-sensitive). Alignments were filtered (≥95% nucleotide identity) by using pysam to ensure species-level resolution and downstream SNV analysis. MAG abundances were calculated as transcripts per million (TPM), normalizing genome coverage by total sequenced bases per sample. Microbial diversity was evaluated at both the alpha and beta levels:

- Alpha diversity (richness and evenness):

- MAG richness was quantified per sample using genome breadth detection thresholds specific to each domain (>5% for eukMAG, >10% for proMAG, >30% for pMAG and vMAG) with minimum read depth (≥1×) to reflect domain-specific variation in genome size.
- Evenness was quantified using the Shannon diversity index, calculated from the relative abundance using the scikit-bio Python package.
- - Beta diversity was assessed using robust Aitchison distances, computed with the vegdist function in the vegan R package. Principal coordinates analysis was performed using cmdscale to visualize community-level dissimilarities across samples.

Human-associated viruses (vMAGs) classified into classes Papovaviricetes, Revtraviricetes, or Pokkesviricetes were identified using geNomad^21^. Samples containing these viruses above detection thresholds were defined as “host-infected” and used as explanatory variables in diversity analyses.

To identify factors shaping microbial community variation, PERMANOVA was performed using the adonis2 function in vegan. Metadata variables—country, skin type, gender, disease status, and host infected—were each tested independently, then cumulatively added in descending order of marginal R² to estimate their additive explanatory power. Pairwise comparisons within categorical variables (e.g., between countries or skin types) were conducted using the pairwiseAdonis extension.

### Human genomics imputation and association analysis

We performed host genomic variant detection and genome-wide association analysis on 1,756 metagenomic samples using an ultra-low coverage imputation approach. Raw sequencing reads were mapped to the human reference genome (GRCh38.p14) using BWA-MEM^37^ (v0.7.17). For individuals with multiple samples (as detailed in Supplementary Table 4), BAM files were merged using the merge function in Samtools^11^, to consolidate host-derived reads across time points or body sites. The merged BAM files were processed using the GATK^38^ (v4.5.0.0) best practices pipeline, consisting of the following sequential steps: MarkDuplicates, BaseRecalibrator, ApplyBQSR, and HaplotypeCaller. Variant calls were subsequently partitioned into genomic chunks suitable for imputation.

Genotype imputation and phasing were performed using GLIMPSE2^39^ (v.2.0) with the 1000 Genomes Project reference panel, retaining samples with average genome coverage >0.1%. After filtering, 327 individuals from four countries remained (76 individuals excluded due to low coverage). Imputed chromosome-level VCF files were merged to create a single autosomal variant dataset per individual.

Variants were filtered using BCFtools^40^ (v.1.17) and VCFtools^41^ (v.0.1.16), retaining high-confidence, biallelic SNPs (minor allele frequency >5%; missing genotype calls <10%), resulting in 5.85 million SNPs.

We conducted GWAS using PLINK2^42^ (v2.00a5). To account for population stratification, we calculated the top ten principal components (PCs) using --pca function. The first four PCs explained 38.71% of the total genetic variance and clearly distinguished clusters by ethnicity; these were thus included as covariates in all association models.

To address redundancy arising from multiple metagenomic samples per individual (different body sites and time points), we performed one hundred iterations of random subsampling. In each iteration, a single representative sample per individual was randomly selected, using an incremented random seed (from 1). For USA samples, analyses were restricted to four skin sites (Al, Ck, Gb, and Ac) to harmonize anatomical site distribution across countries. MAGs were included if present in >20% of 327 individuals and their abundances were transformed using the centered log-ratio (CLR).

A generalized linear model was applied in PLINK2^42^, treating each MAG abundance as the phenotype. Covariates included the first four PCs, country, gender, and disease status. Skin type was excluded due to multicollinearity (variance inflation), particularly its confounding with country. SNP–MAG associations were tested in each of the 100 iterations; SNPs were considered consistently significant if they achieved a *P* value less than 5×10LL in at least 90 of the 100 subsampling tests. Across analyses, 22 MAGs consistently showed significant associations, prompting a Bonferroni-adjusted study-wide significance threshold of 2.27×10LL (5×10LL divided by 22 MAGs). Furthermore, to quantify the strength of genotype–MAG associations, we fitted a mixed-effects model including individual ID as a random intercept. The full model was specified as:

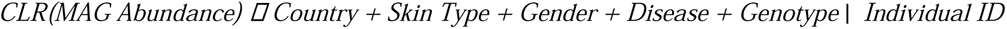

We calculated a McFadden-like pseudo-R² by comparing this full model with a reduced model excluding the genotype term, enabling quantification of the marginal contribution of genotype to microbial abundance variation.

Manhattan plots were generated using GWASlab, based on the median P value across the 100 iterations. Significant SNPs were annotated with rsIDs and nearest gene identifiers. To evaluate biological relevance to skin tissue, gene expression was queried using the Human Protein Atlas database^43^ (https://www.proteinatlas.org/humanproteome/tissue/skin), specifically considering genes expressed in “skin” categories and genes with skin-enriched RNA expression profiles.

### Human population structure and microbial composition comparison

To investigate the relationship between host genetic structure and microbial community composition, we compared human population stratification with individual-level MAG profiles. Procrustes analysis was employed to assess the concordance between the first four PCs of host genotype data and microbial composition ordination. Microbial dissimilarities were calculated using the robust Aitchison distance, and Mantel tests were conducted to evaluate the correlation between genetic and microbial distance matrices. These analyses were repeated across one hundred independents random subsampling of MAG abundance profiles to ensure robustness against sampling variability.

To account for potential geographic confounding, partial Mantel tests were conducted with geographic distance as a covariate. Geographic distances between individuals were computed using the Haversine formula, based on the latitude and longitude of the sampling cities in the four included studies. This approach allowed us to distinguish the independent contribution of host genetic variation to microbiome structure.

### Microbial SNV analysis

Since reliable SNV calling requires sufficient read depth and genome coverage^44^, we first selected sample-MAG pairs with >70% genome breadth with 10-fold coverage. To enable cross-country comparisons, we included only those MAGs that were present in more than 10 samples. SNV analysis was performed using Samtools’ mpileup module^11^ with a mapping quality filter (-C 50) to align bases to each reference. Each SNV was defined as a single nucleotide polymorphism if the alternative allele was supported by a frequency greater than 1% and at least four reads. The microdiversity (nucleotide diversity, π) of each genome was calculated using an equation from Schloissnig et al^45^.

### Microbial pN/pS Estimation

Following the construction of the eSMGC and the SNV analysis pipeline described above, we calculated pN/pS to evaluate selective pressures on coding sequences. Briefly, for each coding gene within the eSMGC MAGs, codons were extracted in frame and assessed for potential singleLnucleotide changes. Each codon’s expected nonsynonymous (eN) and synonymous (eS) sites were determined by enumerating all singleLbase substitutions (3 positions × 3 possible mutations) and classifying each substitution as aminoLacidLchanging (nonsynonymous) or silent (synonymous). Under the assumption of uniform mutation probabilities, the total count of nonsynonymous substitutions for a codon *C*, nonsyn(*C*), was scaled by 1/3 to yield eN(*C*), with eS(*C*) = 3 - eN(*C*).

All observed SNVs passing the aforementioned coverage (≥10×) and alleleLfrequency (>1% with ≥4 reads) filters were mapped back to their respective codons. Substitutions altering the amino acid were summed as nonsynonymous variant frequencies (fN), whereas silent substitutions were summed as synonymous variant frequencies (fS). Per gene, pN and pS were calculated as

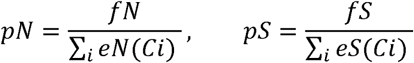

where the sums are over all codons *Ci* in the gene. The final ratio pN/pS was then used to compare rates of nonsynonymous to synonymous variation, providing a populationLlevel analog to dN/dS in metagenomic contexts without haplotype assignment. Genes lacking any qualifying SNVs were assigned zero values for fN and fS, resulting in pN/pS = 0 when no nonsynonymous variation was detected.

### *Cutibacterium acnes* genome-wide association analysis

Variant calling of *C. acnes* MAGs in each sample was generated using a Snippy pipeline (v.4.6.0; https://github.com/tseemann/snippy), which compares individually reconstructed MAG to a reference genome. The most representative *C. acnes* MAGs were selected as references using dRep^25^ from 706 validated MAGs (completeness > 70%, contamination < 5%) obtained through individual and co-assembly, as well as binning methods. Snippy synthesized reads from contigs to call variants, which were then filtered using Freebayes^46^ (v.1.3.6; -p 2 -P 0 -C 2 -F 0.05 --min-coverage 10 --min-repeat-entropy 1.0 -q 13 -m 60 -- strict-vcf) and BCFtools^40^ (FMT/GT=1/1, QUAL>=100, FMT/DP>=10, (FMT/AO)/(FMT/DP)>=0). The filtered variants from the individually binned MAGs (505 in total, excluding 201 co-assembled MAG) were merged into a multi-sample VCF file for further association analysis.

Genome-wide association analysis (GWAS) by geographical effect was conducted using Pyseer^47^ (v1.3.11), with additional covariates fixed effects of sex and skin characteristics from a filtered VCF file. Pyseer^47^ could not calculate the significant variant for more than two phenotypes, so we tested each country (Italy: 10, India: 42, Singapore: 63, China: 149) against the USA sample (241). Significant variants distinguished from the USA samples were identified by likelihood ratio test (lrt) *P* values lower than 5×10LL. Lineage-specific variants were also assessed using the “--distance and --lineage” options in conjunction with a phylogenetic tree generated from 628 MAGs by PhyloPhlAn3^27^. A variant was considered lineage-specific if it significantly correlated with a MDS component (*P* value <0.05) as determined by the Wald test.

### Cutibacterium acnes phage analysis

A total of 126 vMAGs assigned to *Cutibacterium acnes* phage showed clear separation in the draft phylogenetic tree of vMAG with host assignment (Supplementary Fig. 8). To further elucidate their phylogenetic relationships, pan-genome clustering of these vMAGs was conducted using MMseqs2^48^ (v.13.45111) easy-cluster (--min-seq-id 0.5 -c 0.8). This analysis revealed only three clusters comprising more than 75% of the *C. acnes* phage vMAGs. Core gene nucleotide sequences from these clusters were concatenate and aligned using MAFFT^49^ (v.7.520; default option) and trimAl^50^ (v.1.4; -gt 0.9 -cons 60). Phylogenetic trees were subsequently constructed with RAxML-NG^51^ (v1.2.1; --model GTR+G --all).

Phage prevalence was determined for each country separately based on the presence of these vMAGs, and the average relative abundance was visualized using iTOL^52^ (v.6) along with the corresponding phylogenetic tree. Additionally, samples containing *C. acnes* phages were matched with those where *C. acnes* MAGs had been recovered, allowing assessment of associations based on their respective phylogenetic trees. Host-phage coevolution was statistically evaluated using ParaFit^53^ (v5.8.1), based on phylogenetic distances.

To investigate host defense strategies, defense systems within each *C. acnes* genome were predicted using DefenseFinder^54^. Linear regression analyses were conducted to examine the relationship between the number of associated phages and the number of defense systems, controlling for country as a covariate.

### Host virus associated skin microbial diversity

Among the 382 human-associated vMAGs identified, 97.6% belonged to the family *Papillomaviridae*. Phylogenetic analyses were performed using the viral L1 gene, following alignment, trimming, and tree construction methods identical to those described above for *C. acnes* phages. To determine the taxonomic identity and novelty, these vMAGs were compared against 451 complete reference genomes retrieved from the PaVE database^55^ (https://pave.niaid.nih.gov/explore/reference_genomes/human_genomes) using FastANI^56^ (v.1.33; ANI <95%). Taxonomic classifications were further confirmed by aligning L1 genes to reference genomes using BLASTn with previously established thresholds^57^.

To accurately compare microbial diversity between host-virus-infected and uninfected samples, analyses were restricted to samples containing >10 million paired-end reads, with subsampling performed uniformly at 10 million reads per sample. Microbial richness and evenness were calculated as previously described (see “Microbiome profiling”), stratified by microbial domain. Host-associated vMAGs were excluded from viral community diversity metrics.

### Host viral association prediction via machine learning

We constructed a machine learning model to predict host-virus associations based on domain-level diversity metrics, abundances of eSMGC MAGs, and functional gene profiles. Gene abundances quantified using FeatureCounts^58^ (v1.6.0) were summarized by KEGG orthologs and normalized (TPM). Feature selection was performed using the BorutaPy algorithm, retaining the top 50 informative features. To prevent data leakage across different skin sites within the same individual, model validation was performed using five-fold cross-validation stratified by individual, implemented with StratifiedGroupKFold from scikit-learn.

An XGBoost classifier was utilized, with hyperparameters optimized via GridSearchCV across the following parameter space: max_depth: [3, 5, 10]; n_estimators: [200, 250]; learning_rate: [0.1]; min_child_weight: [1, 5, 10]; gamma: [0.5, 1, 1.5, 2]; subsample: [0.6, 0.8, 1.0]; and colsample_bytree: [0.6, 0.8, 1.0]. Model performance was evaluated using AUROC, accuracy, recall, and precision metrics computed via scikit-learn.

Additionally, significant microbiome associations (MAGs and their functional gene profiles) with host viral infection were identified using MaAsLin2^59^. A mixed-effects model was applied with host viral infection status as the dependent variable, incorporating gender and country as fixed-effect covariates, and adjusting for individual ID and skin type as random effects. Furthermore, KEGG orthologs significantly associated with host virus presence (Benjamini–Hochberg-adj. *P* < 0.1) were utilized to calculate pathway enrichment using MinPath^60^ (v.1.6).

## Data availability

The genome database of the expanded Skin Microbial Genome Collection (eSMGC) is available on Figshare: https://figshare.com/s/c7372de026daebb610fb.

## Acknowledgments

H.J.S. acknowledges the support of Korea National Institute of Health (KNIH) research project (project No:2024ER211600); Korea Environment Industry & Technology Institute (KEITI) through the Core Technology Development Project for Environmental Disease Prevention and Management, funded by the Korea Ministry of Environment (grant number: RS-2022-KE002048). C.Q. acknowledges the support of the Biotechnology and Biological Sciences Research Council (BBSRC), part of UK Research and Innovation; Earlham Institute Strategic Programme Grant (Decoding Biodiversity) BBX011089/1 and its constituent work package BBS/E/ER/230002C; the Core Strategic Programme Grant (Genomes to Food Security) BB/CSP1720/1 and its constituent work packages BBS/E/T/000PR9818 and BBS/E/T/000PR9817; and the Core Capability Grant BB/CCG2220/1.

## Author contributions

Methodology and Visualization H.J.S.; Writing – Original Draft, H.J.S.; Writing – Review and Editing,

H.J.S. & C.Q.; Supervision, H.J.S & C.Q.

## Competing interests

All authors declare no financial or non-financial competing interests.

## Notes

### Competing Interest Statement

The authors have declared no competing interest.

