## Supplementary information for "Geographic Variation and Host Genetics Shape the Human Skin Microbiome"

Supplementary Figures 1 to 13

Supplementary Tables 1 to 4

### Supplementary Figures and Legends

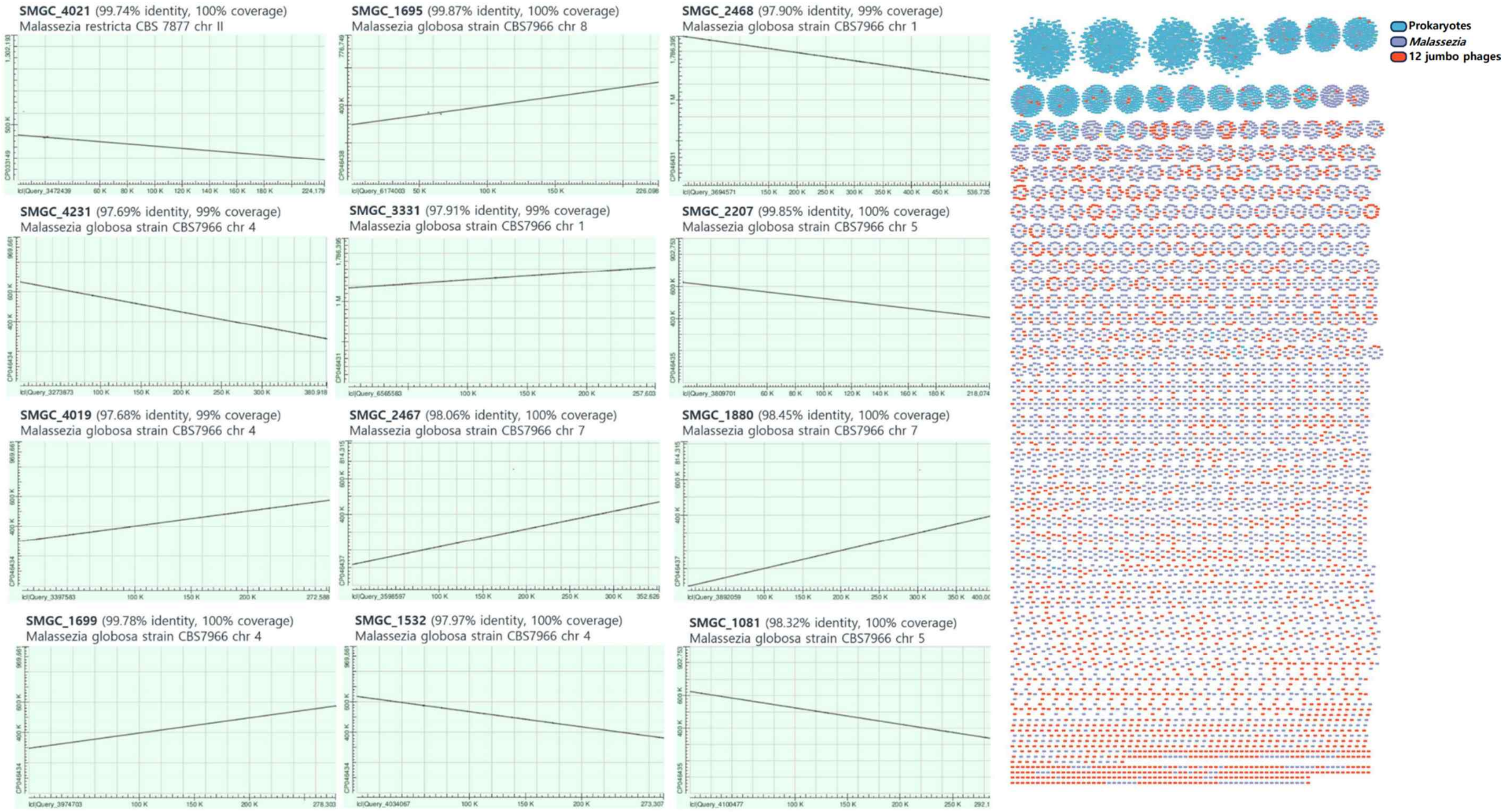


**Supplementary Figure 1. Twelve jumbo phages sequences are part of *Malassezia* genome sequences.** Left panel: BLASTn alignments between the twelve jumbo phage genomes and chromosomes of *Malassezia globosa* strains. The x-axis represents the nucleotide positions of the jumbo phage sequences, while the y-axis indicates the aligned regions of *M. globosa* chromosomes. Right panel: Gene similarity network illustrating relationships among genes from proMAGs (prokaryotic metagenome-assembled genomes), eukMAGs (eukaryotic MAGs, specifically *Malassezia*), and the twelve jumbo phages, derived from the expanded Skin Microbial Genome Collection (eSMGC). Genes from proMAGs, eukMAGs, and jumbo phages are colored blue, purple, and red, respectively.

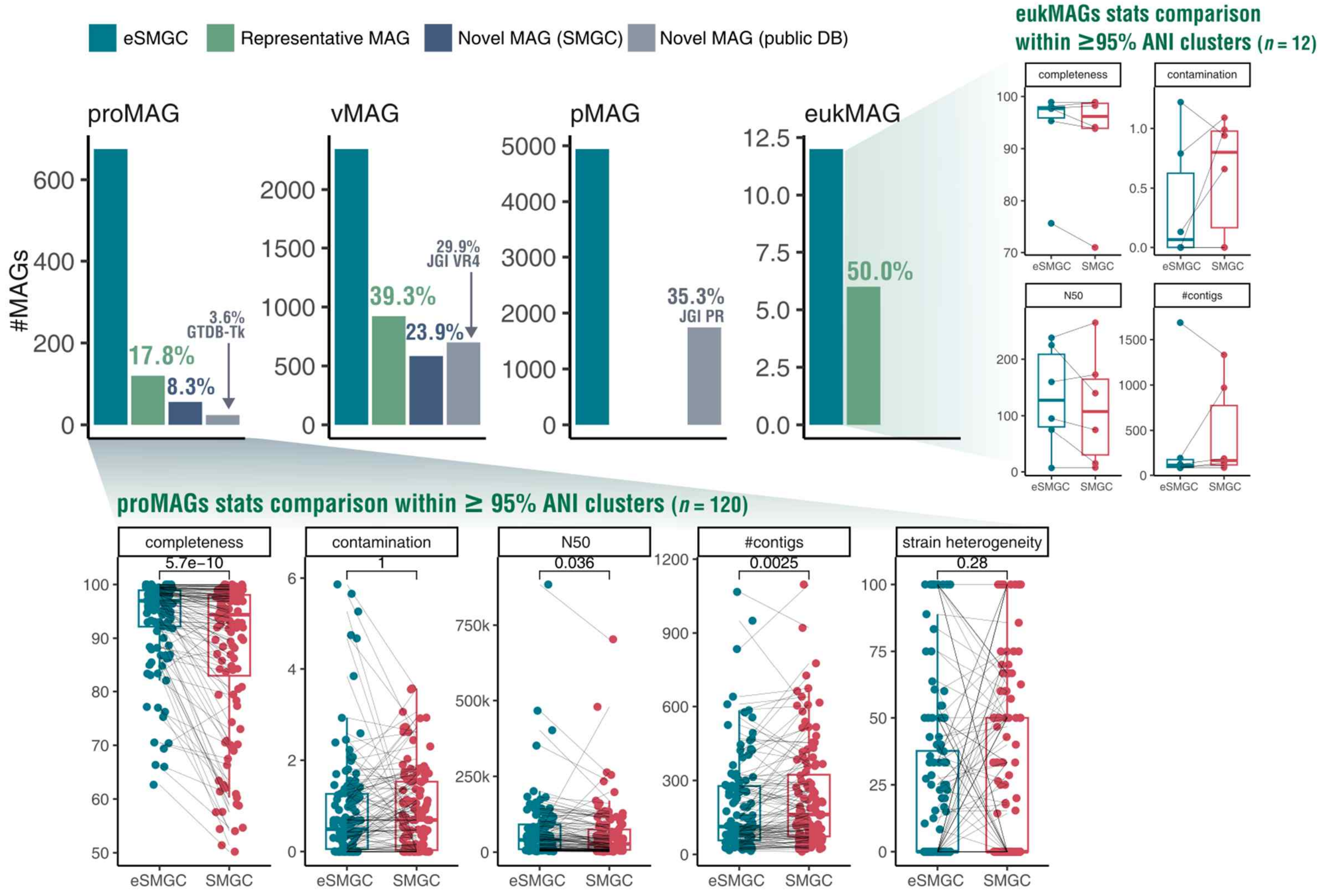


**Supplementary Figure 2. Comparison of the expanded Skin Microbial Genome Collection (eSMGC) with publicly available genome collections.** Bar plots shows the number representative or novel MAGs identified in eSMGC, in comparison to those found in public genome databases. Box plots depict paired comparisons of genome quality metrics for MAGs within ≥95% average nucleotide identity (ANI) clusters, calculated using CheckM for proMAGs and EukCC for eukMAGs. Statistical significance is indicated for each comparison.

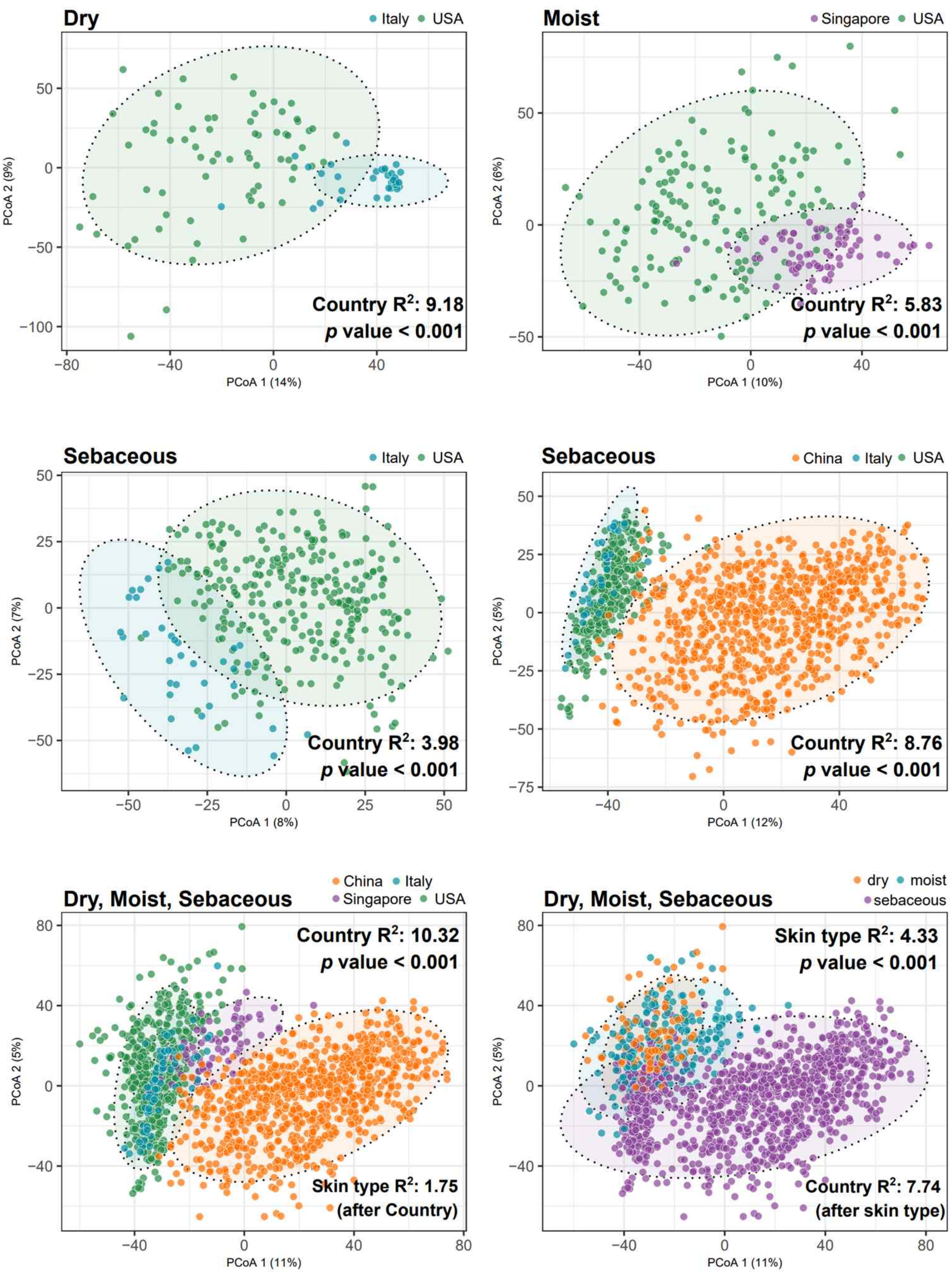


**Supplementary Figure 3. Skin microbial structure stratified by skin type.** Principal coordinate analysis (PCoA) of skin microbiome composition based on Robust Aitchison distances. Each plot displays the effect of country on microbial community structure within specific skin types (dry, moist, sebaceous) or across all types. Variance explained by country and skin type is indicated by the $R^2$ values, as determined by multivariate ANOVA.

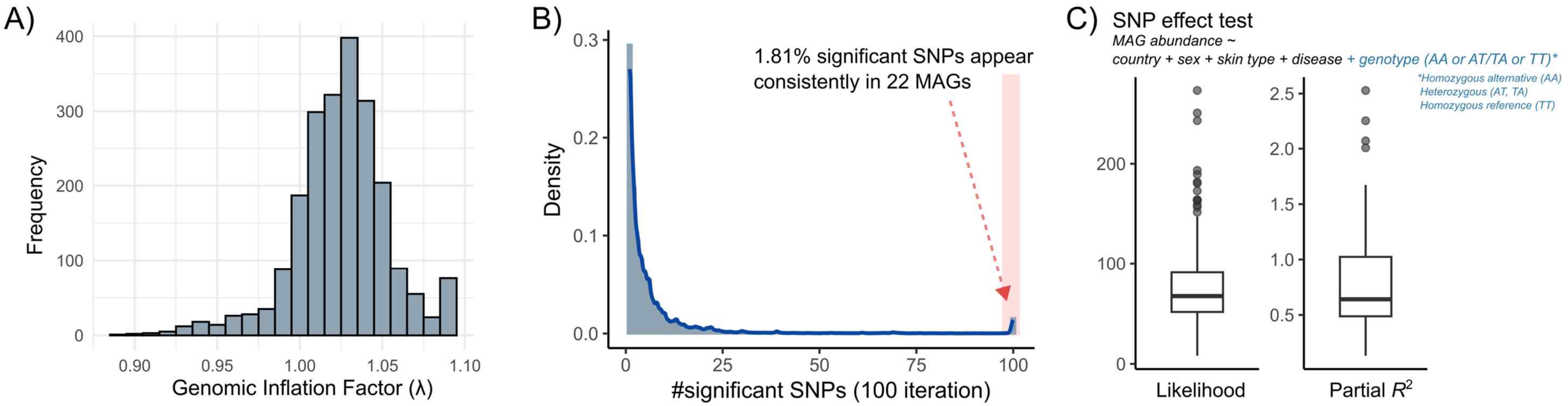


**Supplementary Figure 4. Statistics of genome-wide association analysis of human genetic and skin microbiome abundance.** A) Distribution of the genomic inflation factor (λ) across 100 iterative genome-wide association tests, each involving human genotypes and the relative abundances of 137 microbial species. B) Density distribution of the number of SNPs that reached nominal genome-wide significance ($P$ value $< 5\times10^{-8}$) across 100 replicate tests. A subset of SNPs (1.81%) was consistently significant in association with 22 microbial MAGs. C) Results of mixed-effect models testing the association between consistently significant SNPs and MAG abundance, adjusting for country, sex, skin type, and disease status. The effect of genotype (homozygous alternative [AA], heterozygous [AT or TA], and homozygous reference [TT]) was evaluated after controlling for covariates. Model fit was assessed using likelihood, and explanatory power was quantified by McFadden's partial $R^2$, calculated as the difference in $R^2$ between models with and without the genotype term.

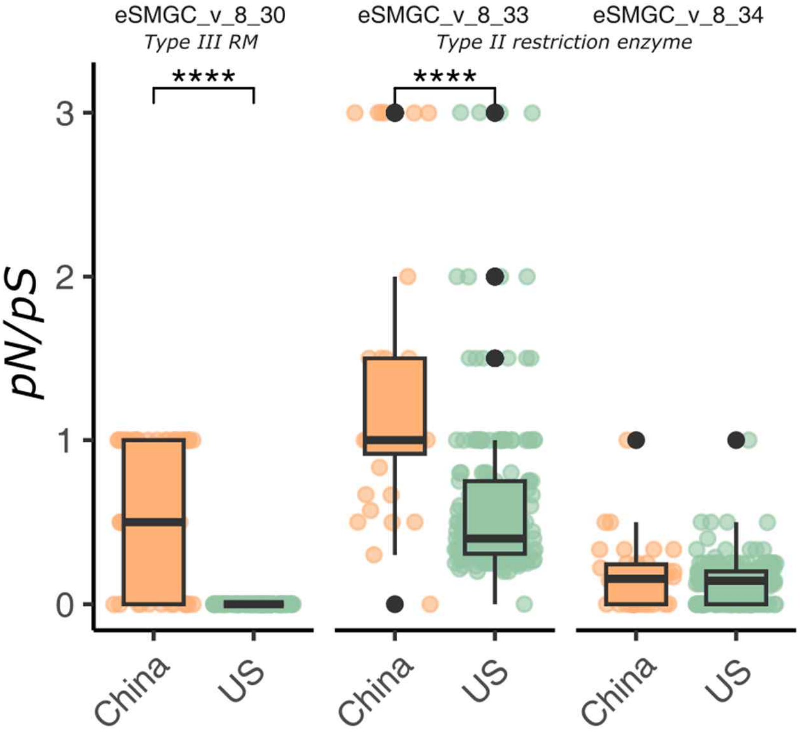


**Supplementary Figure 5. Polymorphism in restriction-modification system genes of *Cutibacterium acnes* prophage (eSMGC_v_8) between China and USA samples.** Boxplots show the ratio of nonsynonymous to synonymous polymorphisms (pN/pS) for three restriction-modification system genes (eSMGC_v_8_30, eSMGC_v_8_33, eSMGC_v_8_34) in *C. acnes* prophages from Chinese and U.S. samples. eSMGC_v_8_30 (Type III RM system) and eSMGC_v_8_33 (Type II restriction enzyme) exhibit significantly elevated pN/pS ratios in the Chinese population compared to the USA population (Wilcoxon rank-sum test, **** $P < 0.0001$), indicating possible differences in selective pressure.

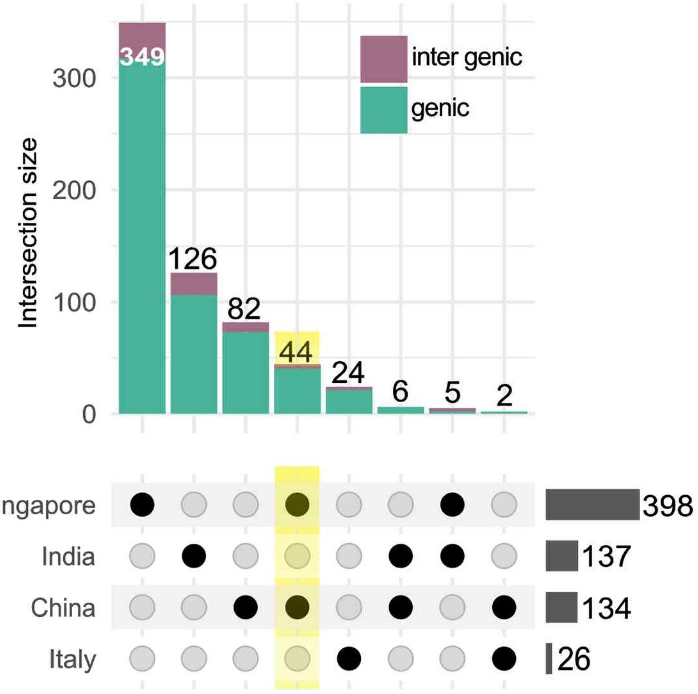


**Supplementary Figure 6. Lineage-independent SNVs in *Cutibacterium acnes* across geographic populations.** Upset plot summarizing lineage-independent SNVs in *C. acnes* strains from Singapore, India, China, and Italy, compared to strains of U.S. origin. Bars indicate the size of SNV intersections across countries, with genic and intergenic variants distinguished by color. The most frequent shared SNVs occur between pairs of countries, with the intersection of SNVs shared among China and Singapore highlighted.

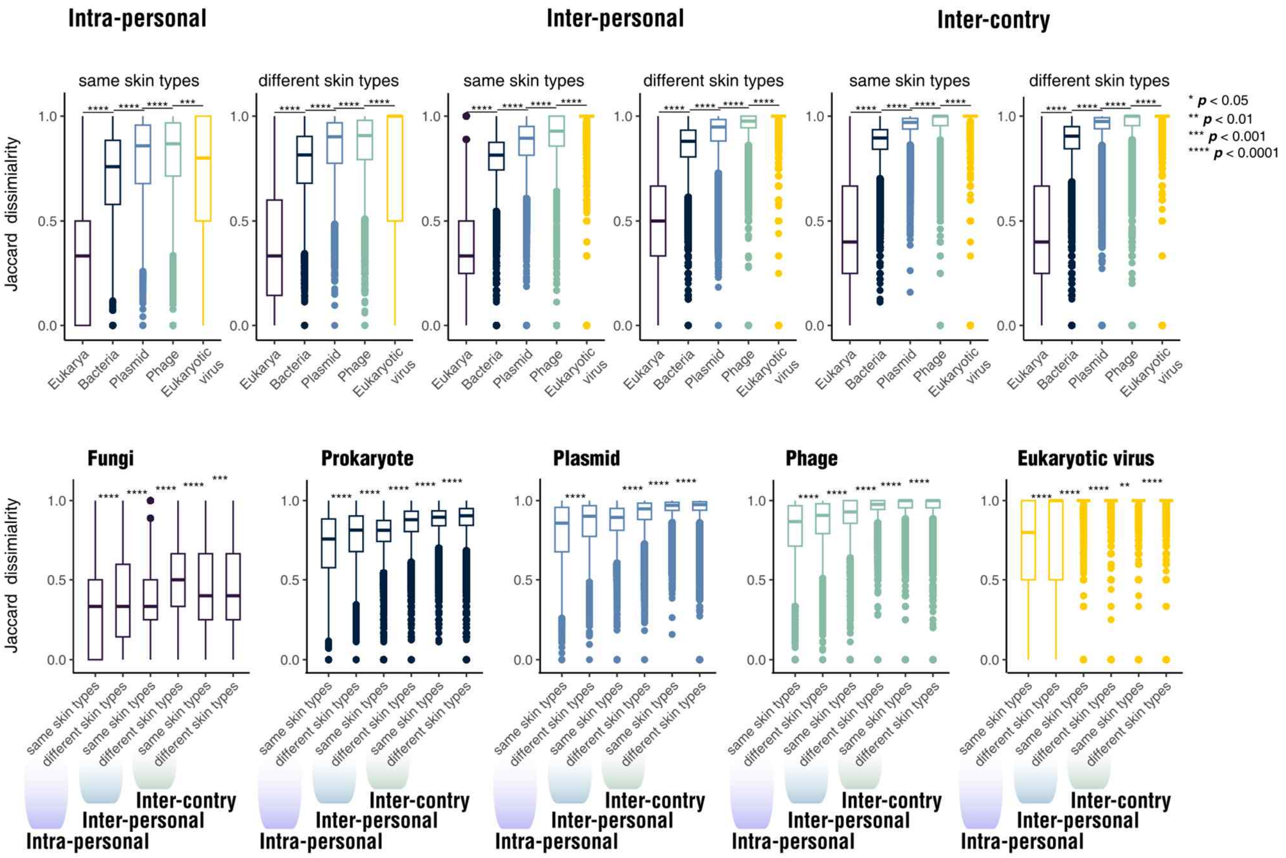


**Supplementary Figure 7. Multi-kingdom compositional stability of the skin microbiome across individuals, skin types, and geographic locations.** Top panel: Boxplots present Jaccard dissimilarity of skin microbial communities across three spatial levels (intra-personal, inter-personal, and inter-country), further stratified by skin site similarity (same or different skin types). Microbial taxa are grouped into eukaryotes, prokaryotes, mobilome elements (plasmids and phages), and eukaryotic viruses. Bottom panel: Jaccard dissimilarity of skin microbiome composition is compared across spatial levels for each microbial group. All domains show increased dissimilarity from individual to country level. Statistical comparisons were performed using Wilcoxon rank-sum tests and significance denoted as: $P < 0.05$ (*), $P < 0.01$ (**), $P < 0.001$ (***), and $P < 0.0001$ (****).

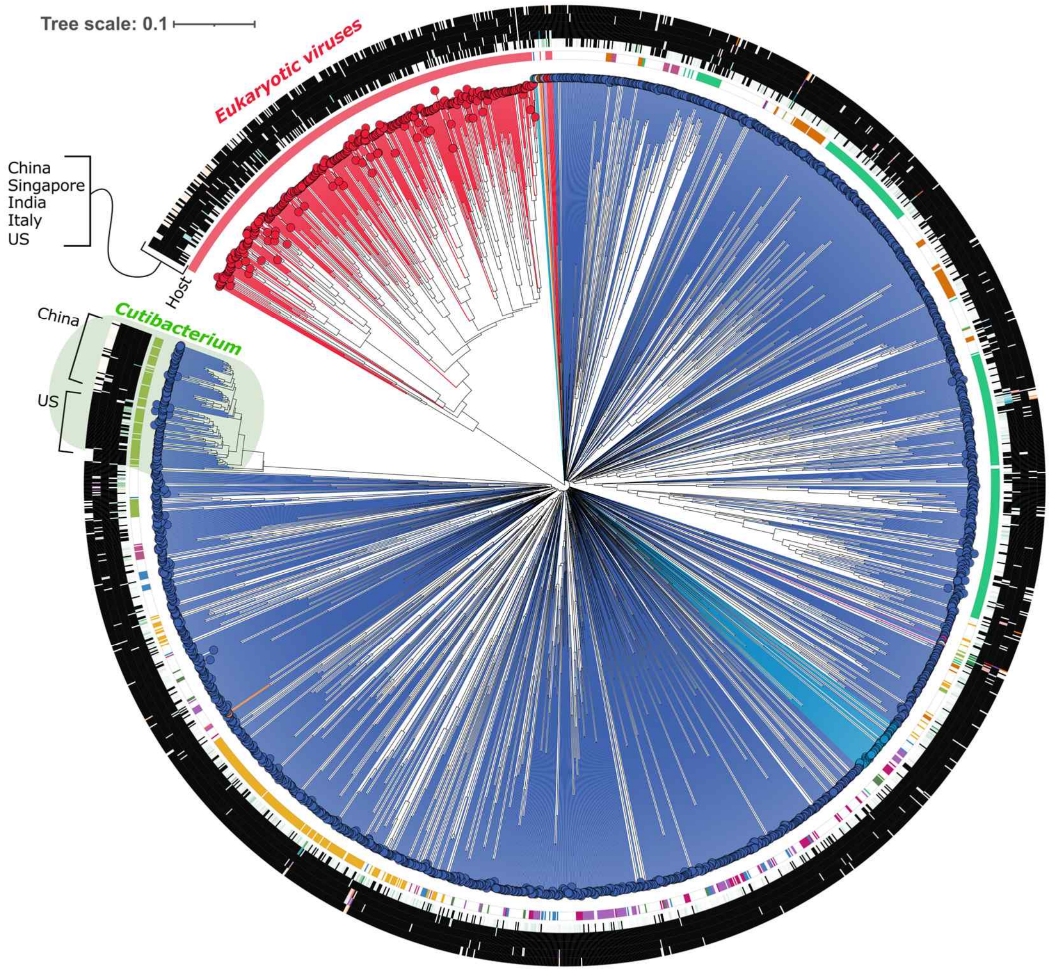


**Supplementary Figure 8. Draft phylogeny of phage and eukaryotic virus genomes recovered from eSMGC.** A phylogenetic tree of vMAGs was constructed using ViPTree based on genome-wide similarity. Branch colors indicate viral families as classified by geNomad. The inner ring denotes predicted host taxonomy, as inferred by iPHoP. The outermost ring shows the proportion of samples in which each vMAG was detected, stratified by country of origin (China, Singapore, India, Italy, and the USA). Tree scale indicates branch length based on genome similarity.

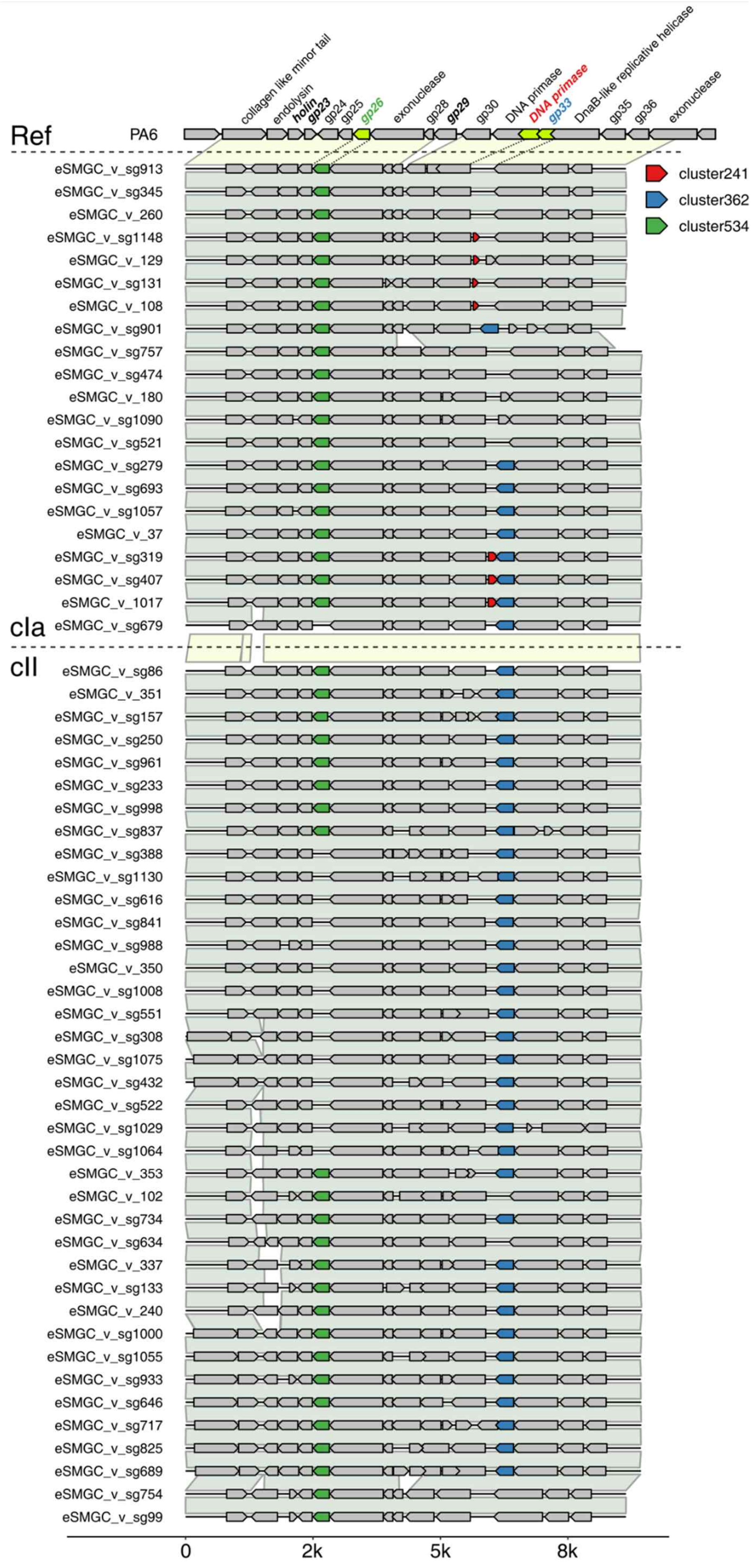


**Supplementary Figure 9. Whole genome alignments of *Cutibacterium acnes* phages across phylogenetic clades.** Whole-genome alignments and synteny plots illustrate genetic variation among *C. acnes* phages, grouped by clades as defined in Fig. 5A. Gene annotations are based on the reference *Cutibacterium* phage PA6 genome. Colored gene clusters correspond to those distinguishing the clades shown in Fig. 5B, highlighting conserved and variable regions across the genomes.

.

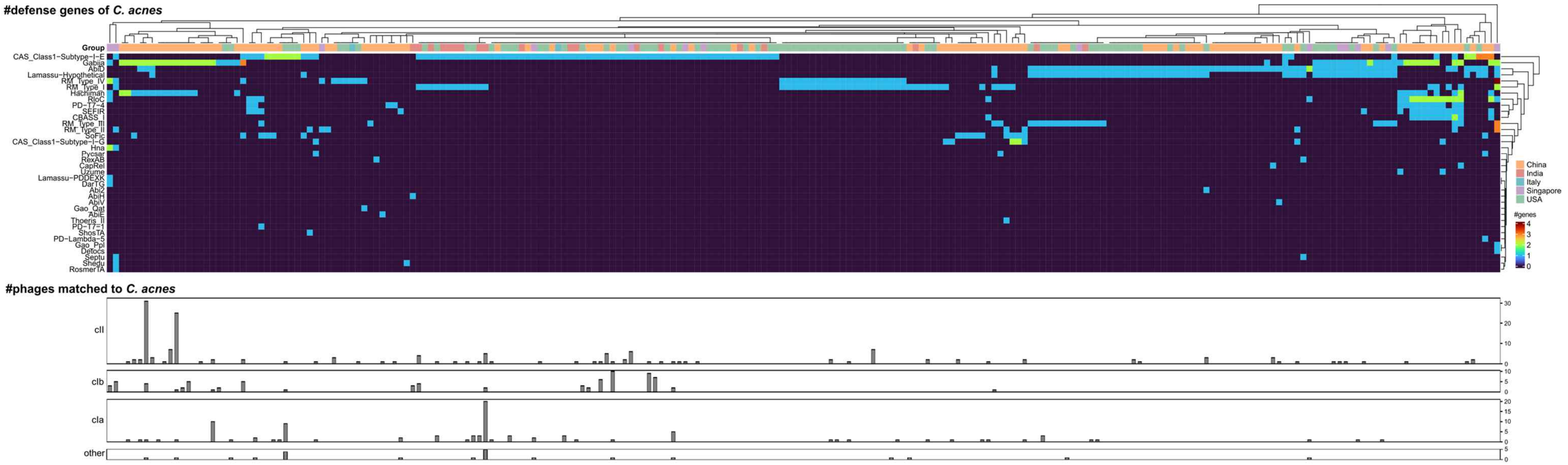


**Supplementary Figure 10. Diverse defense system patterns in *Cutibacterium acnes* show less association with their co-existing phage clusters.** Top panel) Heatmap showing the number and type of *C. acnes* defense system genes recovered per sample, including CRISPR-Cas, restriction-modification, Lamassu, and other systems. Gene counts are color-coded, and samples are annotated by country of origin. Bottom panel) Bar plots indicating the number of *C. acnes*-associated phages detected in each sample, stratified by phage clade (cla, clb, clc, and other). The x-axis represents individual metagenomic samples.

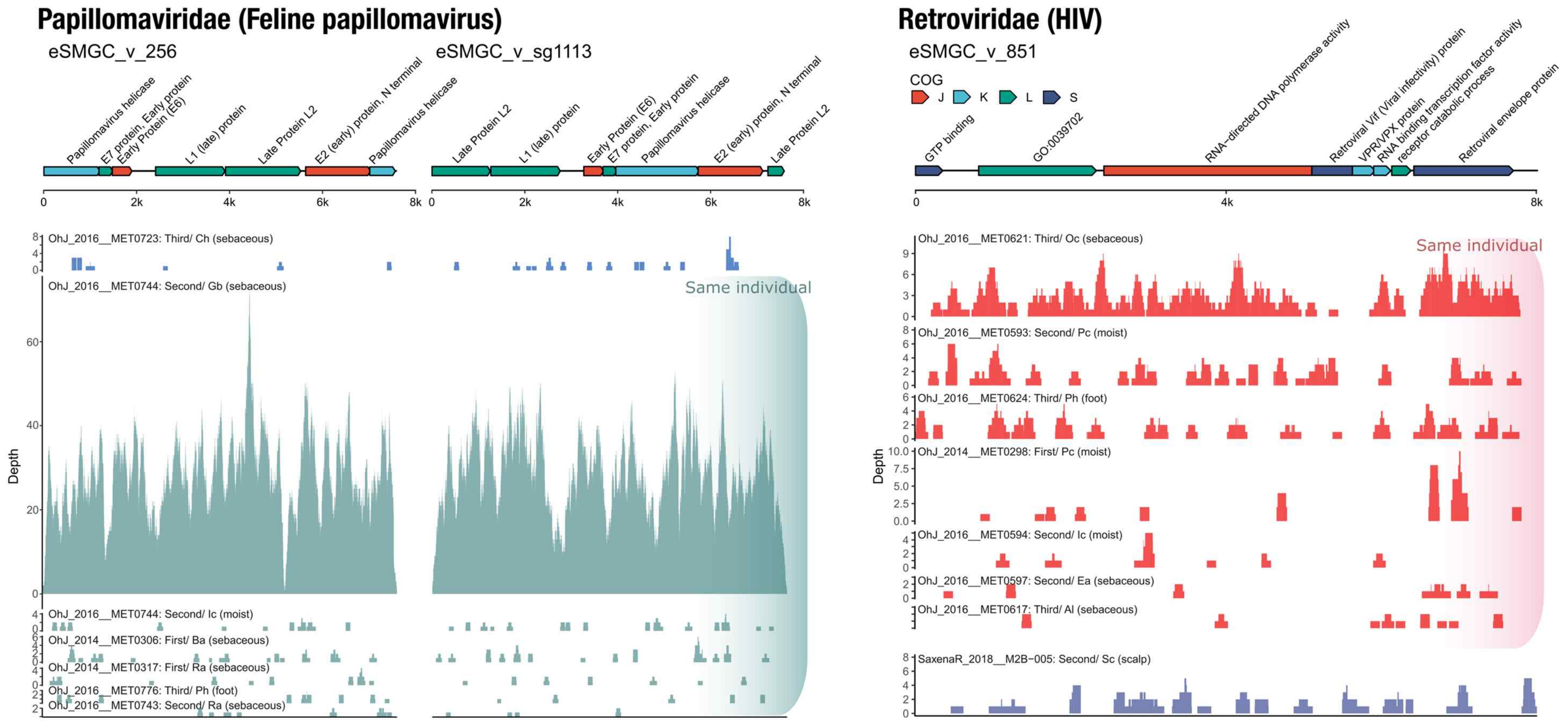


**Supplementary Figure 11**. **Persistence of eukaryotic viral sequences within individuals across timepoints and sampling sites.** Sequences corresponding to feline papillomavirus (Papillomaviridae; left) and human immunodeficiency virus (Retroviridae; right) were consistently detected in two specific individuals from the Oh et al. study, regardless of the timepoint of collection (first, second, or third visit) or the anatomical sampling site. These findings suggest longitudinal and spatial stability of these viral genomes within the same hosts.

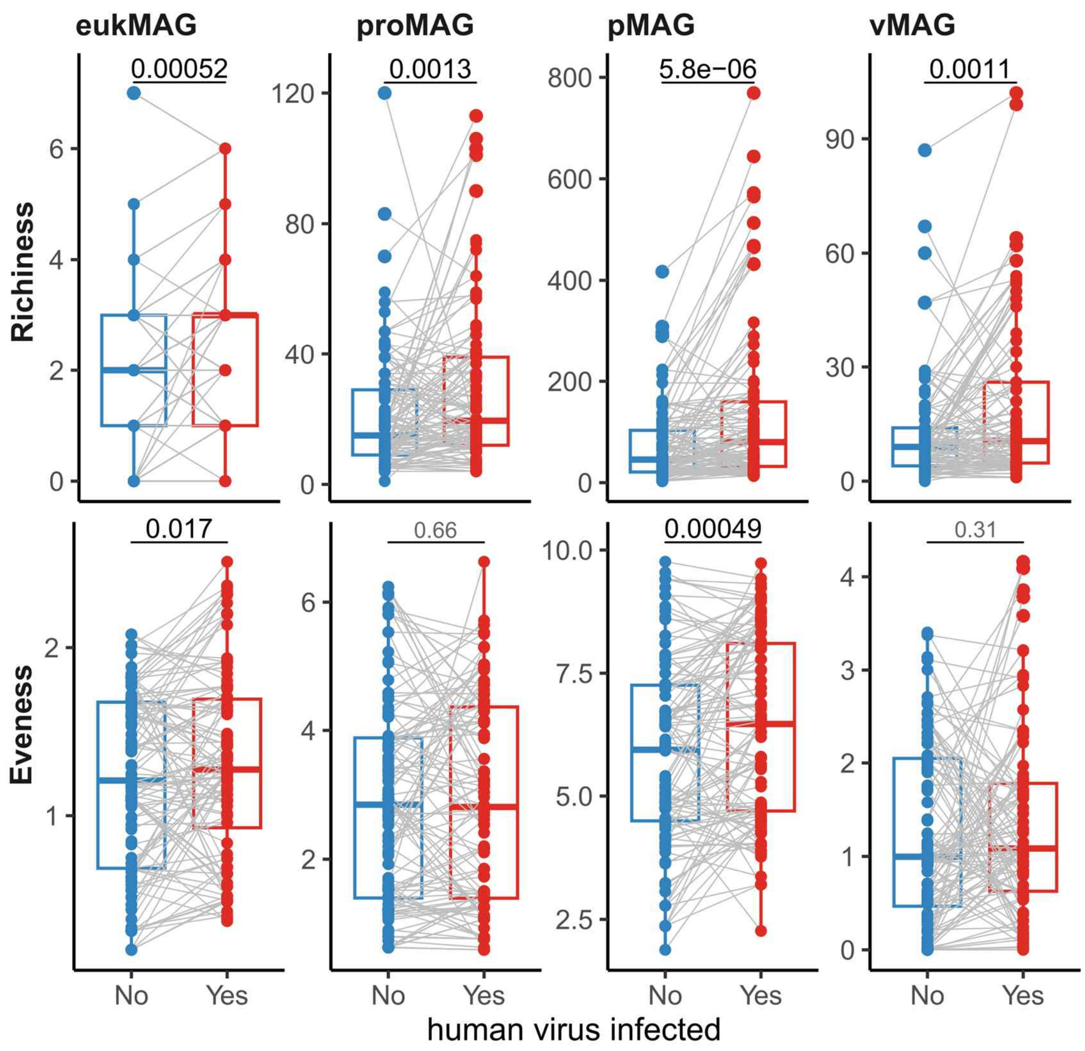


**Supplementary Figure 12. Increased microbial diversity associated with the presence of human viruses on the skin.** Paired samples were analyzed from the same individuals and skin sites, where one sample exhibited the presence of a human virus and the other did not. Boxplots display richness and evenness across four microbial groups: eukMAG, proMAG, pMAG, vMAG (paired Wilcoxon tests).

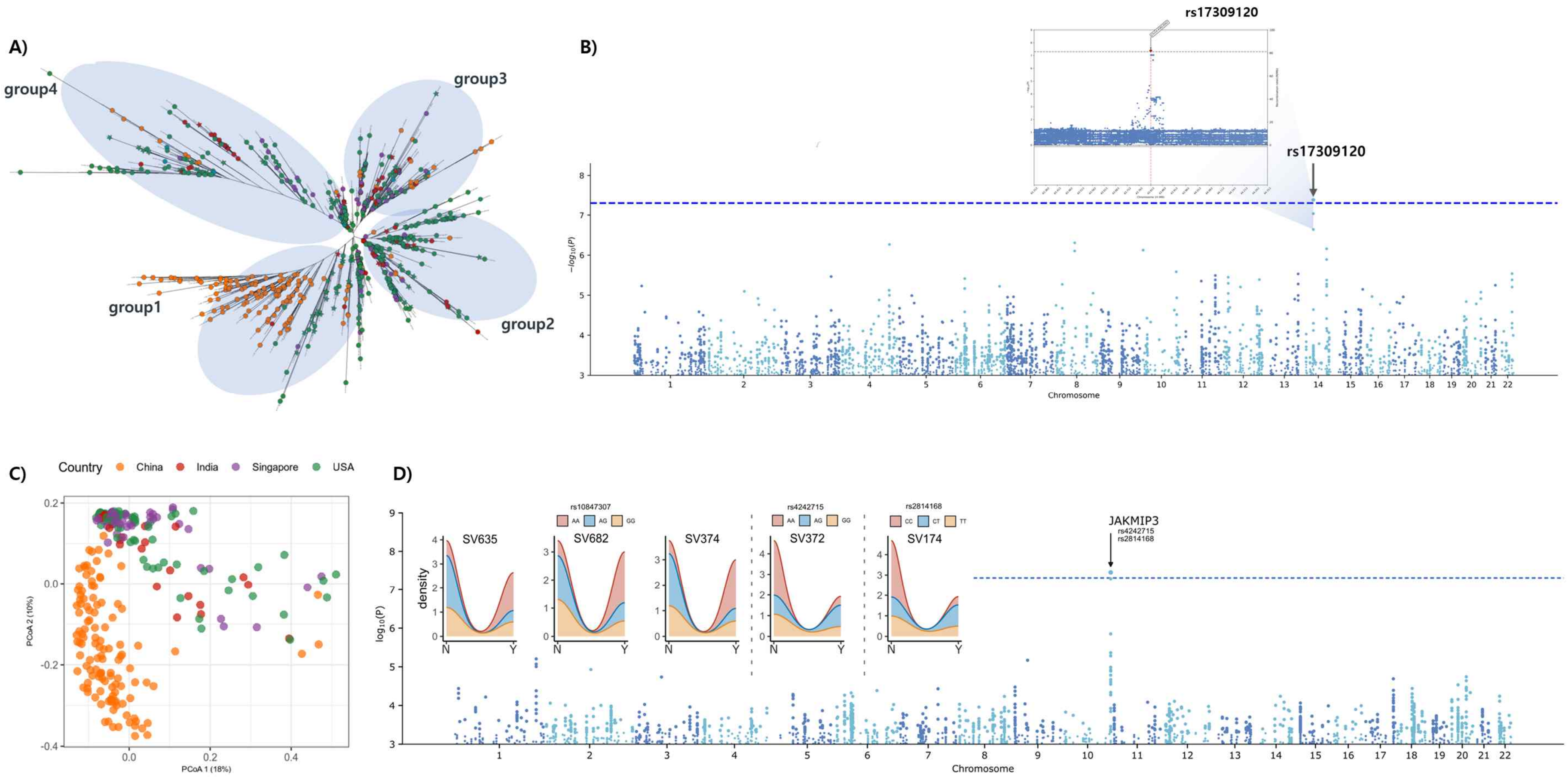


**Supplementary Figure 13. Genome-wide association analysis of *Cutibacterium acnes* genetic and structural variation with host SNPs.** A) Classification of *C. acnes* into four clade groups based on the unrooted phylogenetic tree shown in Fig. 4. B) Manhattan plot depicting a significant association between *C. acnes* clade groups and the host SNP rs17309120. The inset highlights the peak association at chromosome 14. C) Principal coordinate analysis (PCoA) of pan-structural variant (SV) presence among *C. acnes* strains, based on Jaccard dissimilarity. D) Five structural variants (SV635, SV682, SV374, SV372, SV174) were significantly associated with host genotypes. Density plots within panel D show the distribution of SV presence across genotypes for rs10847307, rs4242715 (within JAKMIP3), and rs2814168 (within JAKMIP3). The Manhattan plot in panel D shows a significant association between SV174 and rs2814168. A complete list of genes information of SV is provided in Supplementary Table XX. To avoid redundancy from duplicate host-derived samples, a random subset of strains was used for genome-wide association analysis.

### Supplementary Table Legends

**Supplementary Table 1.** Summary of Consistent Microbial Abundance Changes Across Genotypes in Multiple Countries.

**Supplementary Table 2.** Polymorphism rates of *L. clevelandensis* and *M. chelonae*

**Supplementary Table 3.** *Cutibacterium acnes* (eSMGC_pro_46) pN/pS comparison

**Supplementary Table 4.** Metadata of the skin metagenomic samples, including sample ID, country, sampling site and time point, skin type, and disease status.
